# Efficient enzyme-free isolation of brain-derived extracellular vesicles

**DOI:** 10.1101/2024.01.27.577532

**Authors:** A. Matamoros-Angles, E. Karadjuzovic, B. Mohammadi, F. Song, S. Brenna, B. Siebels, H. Voß, C. Seuring, I. Ferrer, H. Schlüter, M. Kneussel, HC. Altmeppen, M. Schweizer, B. Puig, M. Shafiq, M. Glatzel

## Abstract

Extracellular vesicles (EVs) have gained significant attention as pathology mediators and potential diagnostic tools for neurodegenerative diseases. However, isolation of brain-derived EVs (BDEVs) from tissue remains challenging, often involving enzymatic digestion steps that may compromise the integrity of EV proteins and overall functionality.

Here, we describe that collagenase digestion, commonly used for BDEV isolation, produces undesired protein cleavage of EV-associated proteins in brain tissue homogenates and cell-derived EVs. In order to avoid this effect, we studied the possibility of isolating BDEVs with a reduced amount of collagenase or without any protease. Characterization of the isolated BDEVs revealed their characteristic morphology and size distribution with both approaches. However, we revealed that even minor enzymatic digestion induces ‘artificial’ proteolytic processing in key BDEV markers, such as Flotillin-1, CD81, and the cellular prion protein (PrP^C^), whereas avoiding enzymatic treatment completely preserves their integrity. We found no differences in mRNA and protein content between non-enzymatically and enzymatically isolated BDEVs, suggesting that we are purifying the same BDEV populations with both approaches. Intriguingly, the lack of Golgi marker GM130 signal, often referred to as contamination contamination-negative marker in EV preparations, seems to result from enzymatic digestion rather than from its actual absence in BDEV samples.

Overall, we show that non-enzymatic isolation of EVs from brain tissue is possible and avoids artificial pruning of proteins while achieving a high BDEV yield and purity. This protocol will help to understand the functions of BDEV in a near-physiological setting, thus opening new research approaches.

## Introduction

Extracellular vesicles (EVs) are a heterogenous family of nanosized lipid bilayer-enclosed vesicular structures, which virtually all cell types secrete by evolutionarily conserved processes (Van Niel *et al*, 2018; De Sousa *et al*, 2023). The term EVs includes all currently known subpopulations such as exosomes, microvesicles (MVs), apoptotic EVs, and the more recently described ones such as mitovesicles or migrasomes (Ma *et al*, 2015; Poon *et al*, 2019; D’Acunzo *et al*, 2022). These subpopulations differ in their composition/cargo and biogenesis but are largely indistinguishable by most used isolation methods unless a further specific affinity-based method is used to isolate them from the bulk (Théry *et al*, 2018). EVs contain tissue- and cell-specific cargo composed of proteins (membrane-associated or intraluminal/cytosolic), lipids, and nucleic acids such as DNA (mitochondrial DNA, single- and double-stranded DNA) and RNA (mRNA and diverse non-coding RNAs). By transferring their incorporated and exposed molecules, they elicit biological effects in (targeted) recipient cells (Valadi *et al*, 2007; Van Niel *et al*, 2018; O’Brien *et al*, 2022). Moreover, EVs are naturally surrounded by a corona of attached/associated proteins and other biomolecules that may exert some functions, such as regulating interaction and effects in their target cells (Tóth *et al*, 2021; Wolf *et al*, 2022).

In the central nervous system (CNS), EVs have been highlighted as relevant actors in the complex intercellular communication regulating diverse aspects such as synaptic activity, circuit connectivity, and neuronal differentiation (Antoniou *et al*, 2023; Solana-Balaguer *et al*, 2023; Song *et al*, 2023). However, the transfer of proteins and genetic information between brain cells via EVs is not only relevant in physiology but also in various pathological conditions, such as after ischemic insult (Guitart *et al*, 2016) and in neurodegenerative diseases (ND) (Vassileff *et al*, 2020). A shared feature of ND ((e.g., Alzheimeŕs disease (AD), Parkinsońs disease (PD), and prion-diseases (PrD)) is the aggregation and deposition of characteristic misfolded proteins in distinct areas of the brain, which propagate during the disease course (Hill, 2019; Rastogi *et al*, 2021). Numerous studies have shown that EVs carry and accumulate these pathological proteins (such as amyloid-beta (Aβ) and tau in AD, alpha-synuclein in PD, or transmissible prions in PrD), either inside or at the EV surface/corona, and mediate their propagation (Fevrier et al, 2004; Mattei et al, 2009; Perez-Gonzalez et al, 2012; Ruan et al, 2020; Howitt et al, 2021). Given the EV capability to cross the blood-brain-barrier (BBB) (Ramos-Zaldívar *et al*, 2022), there is a growing interest in defining their cargo in different biofluids, as brain-derived EVs (BDEVs) may provide valuable information about the (patho)physiology of the brain and serve as disease indicators and biomarkers (Vassileff *et al*, 2020). However, despite their promising potential, further research is needed to understand the complete role of BDEVs in ND pathophysiology.

Given the growing interest in and the relevance of isolating EVs from complex tissues such as the brain, the number of protocols is increasing (Vella *et al*, 2017; Hurwitz *et al*, 2018; Brenna *et al*, 2020; Huang *et al*, 2020; Su *et al*, 2021; D’Acunzo *et al*, 2021; Gomes *et al*, 2023; Huang *et al*, 2023; Pait *et al*, 2024). Brain EVs isolation is challenging since BDEVs need to be released from the extracellular matrix (ECM), which consists of an intricate meshwork of (glyco-)proteins, large proteoglycans, and other molecules. In the current EV isolation methods, different approaches, including enzymatic treatment and gentle mechanical dissociation, are employed to address this challenge (Brenna *et al*, 2021). However, the commonly used enzymes like papain and collagenase have already been shown that can alter the protein landscape by introducing unwanted cleavages of proteins present at the EV membrane as the cellular prion protein (PrP^C^) (Brenna *et al*, 2020), which has an important role in the pathophysiology of several ND (Prusiner, 1982; Laurén *et al*, 2009; Urrea *et al*, 2018) and may be relevant in BDEV uptake, trafficking, and function in brain pathologies (Guitart *et al*, 2016; Falker *et al*, 2016; Heisler *et al*, 2018; Brenna *et al*, 2020; D’Arrigo *et al*, 2021). This ‘artificial’ and unintended modification of proteins may be particularly relevant for EV membrane proteins and the EV-corona content and may introduce artifacts that interfere with the study of EV biology (Wolf *et al*, 2022; Tóth *et al*, 2021).

The present study describes the effect of collagenase digestion used during BDEV isolation, which produces undesired protein cleavage of EV-associated proteins in brain tissue homogenates and cell-derived EVs. In order to avoid this effect, we explored the possibility of isolating BDEV with either a reduced amount of collagenase or entirely excluding collagenase from the process. The BDEVs isolated with both approaches were characterized by nanoparticle tracking analysis (NTA), transmission electron microscopy (TEM), western blot, multiplexed mRNA panels, and mass spectrometry, showing no main differences. However, even this minor collagenase treatment still produces the pruning of some EV proteins. We show that the non-enzymatically-based method allows the preservation of membrane proteins while yielding high quantities of BDEVs. This method provides more reliable BDEV samples for diagnosis and research. Intriguingly, our results also show the presence of the Golgi protein GM130 in non-enzymatically isolated BDEVs, therefore questioning its use as a suitable negative marker for EV validation.

## Materials and Methods

### Biological samples and respective ethics statements

#### Human samples

The use of patient specimens for research after completed diagnosis and upon anonymization was in accordance with local ethical standards and regulations at the University Medical Center Hamburg-Eppendorf (UKE, Germany). Autopsy samples of the frontal cortex from non-demented male controls (n = 6) were donated from the brain bank of the Institute of Neuropathology, HUB-ICO-IDIBELL Biobank, Hospitalet de Llobregat, Barcelona, Spain, in compliance with the Spanish biomedical research regulations, including Ley de la Investigación Biomédica 2013 and Real Decreto de Biobancos 2014, and approval of the Ethics Committee of the Bellvitge University Hospital (HUB). Additional information, including age, post-mortem intervals (PMI), neuropathological evaluation, and National Institute of Aging (NIA) staging, are shown in Table 1.

**Table 1:** Frontal cortex autopsy tissues used for different BDEV preparations with collagenase-assisted and collagenase-free methods.

| No. | IDs | Gender | Age | PMI | NIA scoring | Neuropathological findings |
| --- | --- | --- | --- | --- | --- | --- |
| 1 | **PO1** | M | 78 | 2 h 15 min | A0, B0, C0 | Lacunar infarcts |
| 2 | **PO2** | M | 67 | 5 h | A0, B0, C0 | Lacunar infarcts |
| 3 | **PO3** | M | 71 | 12 h | A0, B0, C0 | No neuropathological findings |
| 4 | **PO4** | M | 61 | 3 h 55 min | A0, B0, C0 | Metabolic encephalopathy |
| 5 | **PO5** | M | 85 | 5 h 45 min | A0, B0, C0 | Lacunar infarcts in striatum |
| 6 | **PO6** | M | 64 | 3 h 30 min | A0, B0, C0 | Mesial sclerosis |

\* BDEVs were isolated twice from the PO1, PO2, and PO3 samples for TEM, WB, NTA, proteomics, and nCounter® panels. BDEVs isolated from the PO4, PO5, and PO6 samples were used for the NTA and nCounter® panels. Brain homogenates from PO1, PO2, and PO3 samples were prepared for collagenase digestion and as loading controls for the WBs.

#### Mice

In this study, we performed no experiments with living animals. Breeding and the euthanasia procedure to obtain brain tissue were approved by the ethical research committees of respective national/local authorities: Freie und Hansestadt Hamburg, Behörde für Gesundheit und Verbraucherschutz, Hamburg, Germany (ORG1023). Adult C57BL/6J mice (35-50 weeks old) were purchased from Charles River Laboratories (Paris, France). All experiments were performed following the protocols and guidelines of the Ethical Committee and the directives of the European Union Council 86/609 and 2010/63.

#### Cell lines

Neuro-2a cells (N2a) were ordered via DSMZ (ACC 148) in the Institute of Neuropathology, UKE, Hamburg (ATCC: CCL-131). Cells were maintained in DMEM (Gibco) supplemented with 10% FBS (Invitrogen) at 37°C and 5% CO_2_ atmosphere.

### Homogenization and collagenase digestion of total brain tissue

20% (w/v) homogenates (200 mg/1mL) of C57BL/6J mouse forebrain or human frontal cortex were prepared with a manual dounce homogenizer (with PBS). A control homogenated with PBS and protease inhibitor (PI) cocktail (complete EDTA-free tablet, Roche) was included to assess the role of endogenous proteases. 100 µL of homogenate were treated with (i) collagenase D (Roche) high (40U/mL, 2 mg/mL); (ii) collagenase D low (10U/mL, 0.5 mg/mL); (iii) collagenase type III high (40 U/mL, 1.5 mg/mL) and (iv) collagenase Type III low (10 U/mL, 0.38 mg/mL). One tube was left without enzymes as a control (with and without PI). Samples were then incubated for 30 min at 37°C in a thermomixer at 500 rpm. The reaction was stopped by adding 200 µL of a stop solution of 2× RIPA (100 mM Tris Base, 300 mM NaCl, 2% NP40, 1% Na-Deoxycholate and 0.2% SDS at pH = 8) with 0.1 M EDTA and freshly added PI. Tubes were vortexed and left on ice for 10 min to ensure the activity of inhibitors and detergent, then centrifuged at 12,000*xg* for 10 min at 4°C. The supernatant was collected, 4× sample buffer (SB, 240mM Tris Base, 8% SDS, 40% glycerol, and 0.2% bromophenol blue at pH = 6.8), containing 5% β-mercaptoethanol (to a final concentration of 1×) was added, and the samples were boiled at 95°C for 10 min. The remaining sample was stored at −80°C for further western blotting analysis (see below).

### Isolation of cell culture supernatant EVs

Cells were cultured to approximately 80% confluency in T175 cell culture flasks in DMEM (Gibco) with 10% FBS (Gibco). 18-24 h before EV isolation, cells were washed with PBS, and the medium was changed to the same DMEM but supplemented with 10% exosome-depleted FBS (EXO-FBS-50A-1, System Bioscience). The media was collected on the day of isolation, and differential centrifugation was performed to remove cell debris from the culture medium (200*xg* for 5 min and 7,000*xg* for 20 min). The whole procedure was conducted on ice, and centrifugations were performed at 4°C to avoid degradation. Subsequently, the supernatant was passed through a 0.22 µm filter using the SteriFlip filter system (Merck) and a PVDF 0.1 µm filter (Merck). The flow-through was then transferred into 12 mL polypropylene tubes (Beckman Coulter) and centrifuged at 140,000*xg* (Beckman Coulter Optima L100 XP, SW40Ti, Swing bucket rotor) for 70 min. The final pellet was resuspended in 90 µL PBS containing PI. See Figure 1d for a scheme of the isolation process.

**Figure 1:**
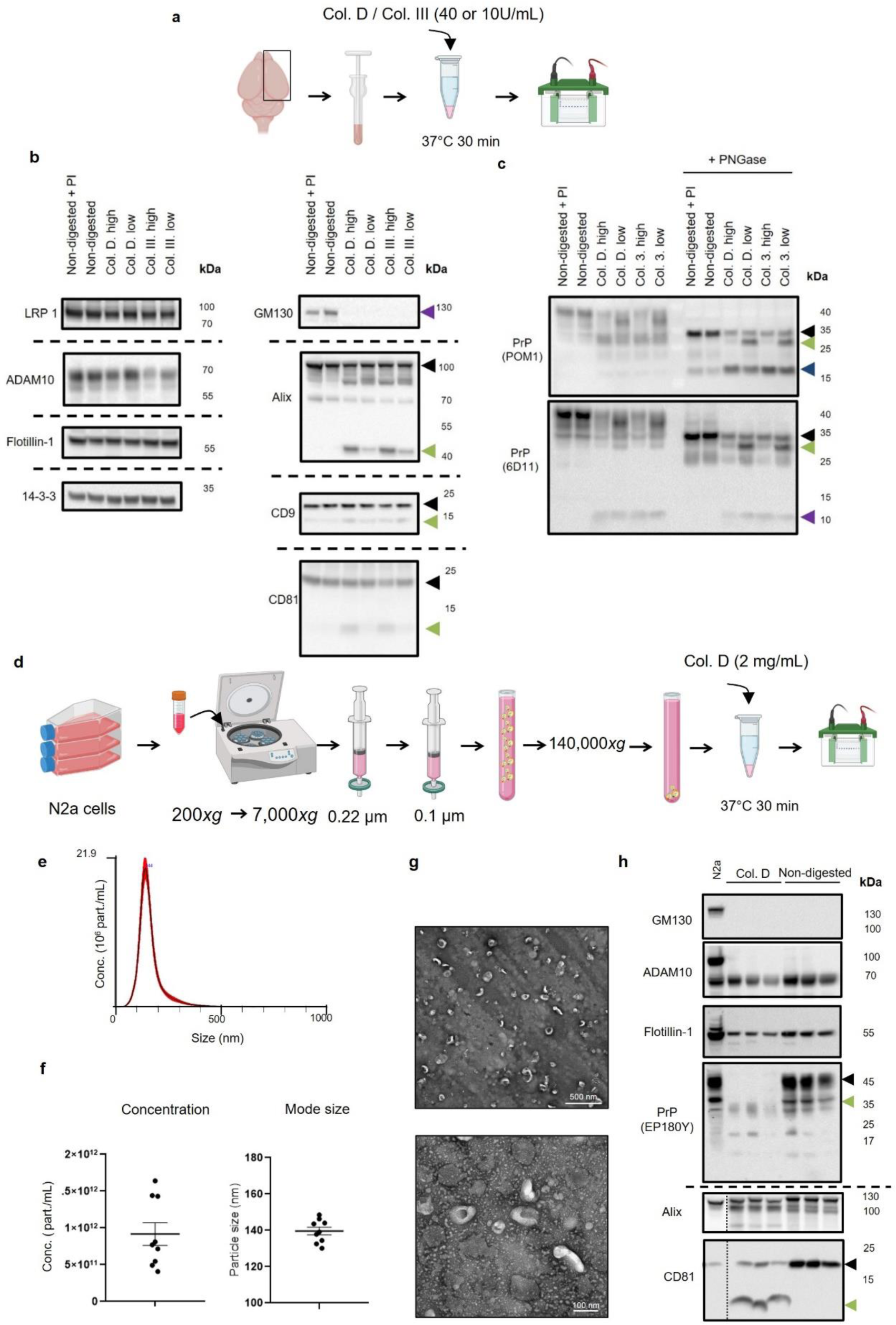
Collagenase digestion induces undesired cleavage of EV-relevant proteins in both brain tissue and cell-derived EVs. (a) Schematic drawing depicting the collagenase treatment in brain tissue. One mouse hemisphere was dissected and homogenized manually in PBS. Subsequently, 100 µL of homogenate underwent treatment with collagenase D and collagenase III at either high (40 U/mL) or low (10 U/mL) concentrations and was then incubated for 30 min at 37°C before being subject to western blotting analysis. (b) Representative western blots of relevant proteins used as EV markers after collagenase treatment. In the left panel, LRP1, Flotillin-1, and 14-3-3 remained unaffected by both collagenase treatments, but ADAM10 amounts were diminished with collagenase III. However, in the right panel, Alix, CD9, and CD81 are visibly cut, producing lower fragments (indicated by green arrowheads). Likewise, GM130 (purple arrowhead) was also found to be digested by collagenase treatments. Non-digested samples with protease inhibitors (PI) and without were used as controls. (c) PrP^C^ patterns after collagenase treatment with and without PNGase digestion were detected by POM1 and 6D11 antibodies. The N-glycan removal shows exacerbated PrP-C1-like (blue arrowhead) and PrP-N1-like (purple arrowhead) fragments, and an artificially generated shorter full-length PrP^C^ (green arrowheads). (d) Schematic drawing depicting the isolation and collagenase treatment of N2a-EVs. EVs were isolated by differential centrifugation and filtration (*n* = 9). Then, the samples were divided into two tubes, each containing 3.0 ×10^6^ EVs, and treated with collagenase D (2 mg/mL, 40 U/mL) or untreated before 30 min incubation at 37°C followed by western blotting analysis. (e) Representative size distribution graph from NTA analysis before treatment. (f) The concentration of particles per mL and mode size analysis (in nm) of N2a-EVs were measured with NTA (*n* = 9). (g) Representative TEM images of N2a-EVs. (h) Representative western blotting of N2a-EVs (*n* = 3 each) treated with collagenase D. N2a cell lysate was used as a loading control. The outer-membrane proteins CD81 and PrP^C^ are altered, but not ADAM10, and the luminal markers Flotillin-1 and Alix. Data are presented as mean ± S.E.M. in (f). Figure created with BioRender.com.

### Isolation of brain-derived EVs (BDEVs)

All brain tissues used here were previously stored at −80°C. During all procedures, protein low-binding tubes (Eppendorf) were used. PBS used in all EV-related experiments was prefiltered through a 0.1 µm filter to avoid impurities. For all the EV resuspensions, a protease inhibitor cocktail at 1× was freshly added to it. Initially, BDEVs were isolated from mouse brain tissues (C57BL/6J, 35-50 weeks old, female) or human brain tissues (see human samples section for details) as follows. The protocol used is based on the one published by Crescitelli *et al*. (Crescitelli et al, 2021), where we added some minor modifications to adapt it to our equipment, reduce the protease activity (i.e., including PI in several steps and excluding or decreasing the amount of collagenase D). The adapted protocol (Protocol A) was as follows:

The tissue dissection and the rest of the procedure were performed on ice to protect samples from degradation. 100-200 mg of frozen brain tissue was minced with a scalpel in a glass petri dish with cold RPMI media (Gibco). 0.55 mL of media was added to each 100 mg of tissue to normalize the media/tissue ratio in all the samples. The mixture was transferred into a 1.5 mL tube (with a cut 1 mL tip), and DNAse I (Roche) was added to a final concentration of 40 U/mL. Collagenase D at low concentration (10U/mL, 0.5 mg/mL) or any collagenase (without, w/o) was added to the samples. Tubes were incubated in a thermomixer at 37°C for 20 min at 500 rpm, and the mixture was pipetted up and down 15 times every 5 min. To stop the endogenous protease activity, RPMI media with 10× concentrated PI (1/10 of the initial volume) was added to the tubes. Subsequently, samples were centrifuged at 300*xg* (5 min), 2,000*xg* (10 min), and 10,000*xg* (10 min). The 10,000*xg* pellet containing large BDEV was not further included in the isolation process and kept as a “10K control” to avoid the larger vesicles and debris. Afterward, the supernatant was transferred into a 5 mL polypropylene tube (Beckman Coulter), and PBS was added to reach a 4.5 mL tube filling and centrifuged at 120,000*xg* for 100 min at 4°C in the L-60 Optima Ultracentrifuge (Beckman Coulter, SW55 TI rotor, Swing bucket rotor). Then, the supernatant was discarded, and the pellet was resuspended with 300 µL PBS (+PI). 240 µL of resuspended BDEVs were mixed with 60% OptiPrep at the bottom of the same tube, while 60 µL were stored at −80°C as the “pre-gradient control” in western blotting analysis. To set up the gradient, 1.3 mL of 30% and 10% OptiPrep (Stemcell) densities (prepared as described in (Crescitelli *et al*, 2021) were layered on top. Finally, 400 µL of PBS was added on top, and the gradient was centrifuged at 185,500*xg* for 120 min at 4°C. Four fractions were collected (F1: 1 mL, F2: 1.3 mL, F3: 0.65 mL, F4: 1.3 mL) into 1.5 mL tubes. BDEVs were expected to mainly concentrate between the 10% and 30% interface. Fractions were then further diluted with PBS, stored at 4°C overnight, and centrifuged at 120,000*xg* for 80 min at 4°C the next day to pellet the EVs and wash out the OptiPrep. After carefully removing the supernatant, the BDEVs were resuspended gently with 60-90 µL of PBS (+PI). Samples were transferred to 1.5 mL tubes, stored at 4°C (if downstream NTA analysis was performed within one week), prepared for further biochemical and imaging analysis (see below), or frozen directly at −80°C.

We also isolated BDEVs from mouse brains (see Supplementary Figure S7) using the protocol previously described by Crescitelli *et al*. (with small modifications to adapt the procedure to our available equipment) (Crescitelli *et al*, 2021) with the collagenase D usual concentration (2 mg/mL). The resulting protocol, Protocol B, shares the same structure as Protocol A besides what is listed below:

i) The tissue was minced with 2 mL of RPMI media, independently of the amount of tissue.
ii) Collagenase D was added to a concentration of 2 mg/mL (40 U/mL).
iii) Tubes were incubated at 37°C for 30 min, not 20 min.
iv) RPMI media with 10× concentrated PI (1/10 of the initial volume) was not added to the tubes.
v) The 10,000*xg* pellet containing large BDEVs was recovered with 200 µL of PBS (+PI).
vi) The 120,000*xg* pellet containing small BDEVs was recovered with 200 µL of PBS (+PI).
vii) A 1:1 mixture of the large and small BDEV (150µL of each) was mixed with 1ml of 60% OptiPrep and included in the gradient.

### Nanoparticle tracking analysis (NTA)

Samples were diluted in PBS depending on estimated EV concentrations. Cell culture-derived EVs were diluted with PBS 1:500 or 1:1,000, while tissue-derived samples were diluted 1:125, 1:250, or 1:500. Before dilution, human BDEVs were treated with 4% paraformaldehyde (PFA) for biosafety reasons. Five videos of 30 seconds duration were recorded (settings: camera level = 16, screen gain = 2) with the NanoSight microscope (LM14, Malvern). The processing function of NanoSight NTA 3.0 software was used to analyze the recordings (settings: detection threshold = 6 and screen gain = 10). PBS (with or without PFA traces mimicking the final concentration in the BDEV samples) was also analyzed as background control and showed almost no signal.

### Negative staining and transmission electron microscopy (TEM)

5 µL of BDEVs were fixed with 16% PFA to a final concentration of 4%. 5 µL of the fixed sample were then added to a formvar/carbon-coated 200 mesh copper grid (#ECF200-Cu-50, Science Services) and left for 20 min in a dry environment. Thereafter, samples were transferred to drops of PBS (3 times for 2 min each). The samples were further stained with 2% methylcellulose-uranyl acetate on ice and looped out on a filter paper after 10 minutes. EVs were analyzed and imaged with a transmission electron microscope (Jeol JEM2100Plus) equipped with a XAROSA CMOS camera (Emsis).

### Collagenase treatment of cell-derived EVs

3×10^10^ particles of N2a-EVs (measured by NTA) were either incubated with or without 2 mg/mL collagenase D at 37°C for 20 min in a thermomixer at 500 rpm. The reaction was stopped by adding 1/8 of RIPA 8× buffer supplemented with an 8× PI to reach a 1× concentration. Samples were left on ice for 10 min for EV lysis and prepared for western blot by adding 4× SB (with 5% β-mercaptoethanol), followed by boiling at 95°C for 10 min.

### Protein quantification

BCA Assay Kit (Pierce) was used according to its manual. The extinction of the bicinchoninic acid (BCA) was measured at 562 nm utilizing the µQuant spectrometer (BioTek). A standard curve was set up using albumin to determine the protein concentration of each sample.

### Protein deglycosylation

Deglycosylation of total brain homogenates was performed according to the PNGase F kit instruction manual (#P0704S, New England Biolabs). 40µg of protein were digested with 1,000 U of PNGase F enzyme.

### Sample preparation for SDS-PAGE

Brain lysates from mouse or human tissues were prepared as a 10% homogenate with RIPA buffer (50 mM Tris Base, 150 mM NaCl, 1% NP40, 0.5% Na-Deoxycholate and 0.1% SDS at pH = 8) with freshly added PI for being used as brain homogenate (BH) controls. N2a cell lysates were prepared using the same buffer. After brain or cell lysis for 10 min on ice, protein lysates were obtained in the supernatants after a 12,000*xg* centrifugation for 10 minutes. EV samples (in PBS, see below) were prepared by adding 8× RIPA buffer (400 mM Tris Base, 1.2 M NaCl, 8% NP40, 4% Na-Deoxycholate and 0.8% SDS at pH = 8) to a final concentration of 1× to not excessively increase the sample volume. Gentle pipetting and a 10-minute incubation were done to guarantee the EV lysis. Finally, in all the cases, 4× SB (containing 5% β-mercaptoethanol) to a final concentration of 1×) was added, and the samples were boiled at 95°C for 10 min.

### SDS-PAGE and western blot analysis

BDEV samples were normalized by the weight of the tissue they were obtained from (BDEV from 25-30 mg of tissue were loaded per lane) and cell-derived EVs by the particle number (measured by NTA). Normalized samples were loaded on pre-casted 4-12% Bis-Tris protein gels (Invitrogen) with a suitable molecular weight marker (10-180 kDa) and loading controls, i.e., either BH or cell lysate. After the electrophoretic separation in MES/SDS buffer (0.5 M MES, 0,5 M Tris base, 1% SDS, and 9.6 mM EDTA), proteins were transferred by wet-blotting with Tris-Glycine buffer (250 mM Tris base, 1.92 M Glycine and 10% methanol) onto a nitrocellulose membrane (BioRad). After completion of the transfer, the total amount of protein was detected using the Revert™ 700 Total Protein Stain Kit (LI-COR) and the Odyssey DLx imaging system (LI-COR) according to the manufacturer’s manual. Subsequently, membranes were blocked for 1 h at room temperature (RT) with 5% non-fat dry milk diluted in TBS-T buffer (100 mM Tris base, 1.4 M NaCl, and 1% Tween-20 at pH = 7.4). Membranes were incubated overnight with primary antibodies at 4°C on a shaking platform. These primary antibodies in a dilution of 1:1,000 were used: 14-3-3 (#9636S, Cell Signaling), ADAM10 (#ab124695, Abcam), Alix (#92880S, Cell Signaling), CD81 human-specific (D3N2D) (#56039S, Cell Signaling) and mouse-specific (D502Q) (#10037S, Cell Signaling), CD9 (#13403S, Cell Signaling or #ab92726, Abcam), Flotillin-1 (#610820, BD Biosciences), Flotillin-2 (#ab96507, Abcam), GM130 (#610822, BD Biosciences or #ab52649, Abcam), Laminin A/C (#SAB4200236, Merck or #ab108595, Abcam), LRP-1 (#ab92544, Abcam), PrP clone POM1 (#MABN2285, Merck), PrP clone EP1802Y (#ab52604, Abcam), PrP clone 6D11 (#808002, BioLegend) and PrP clone 3F4 (#MAB1562, Merck). The next day, corresponding HRP-conjugated secondary antibodies (Anti-Rabbit, #W4011 and Anti-Mouse #W4021, both from Promega) were incubated (1:4,000) for 1 h at RT in 5% non-fat dry milk (in TBS-T buffer). Detection was performed with Pierce ECL Pico or Femto substrate (Thermo Fisher Scientific) using the Chemidoc XRS+ imaging system (BioRad).

### Protein extraction and tryptic digestion of EVs for mass spectrometry analysis

EV samples were diluted 1:1 in 100 mM triethyl ammonium bicarbonate (TEAB), and 1% w/v sodium deoxycholate (NaDoC) and dissolved to a concentration of 70% Acetonitrile (ACN). 2 µg carboxylate modified magnetic beads (GE Healthcare Sera-Mag™) at 1:1 (hydrophilic/hydrophobic) in methanol were added following the SP3-protocol workflow (Hughes *et al*, 2019). Samples were shaken at 1,400 rpm for 18 min at RT and placed on a magnetic rack. The supernatant was removed, and magnetic beads were washed two times with 100% ACN and two times with 70% Ethanol. After resuspension in 50 mM ammonium bicarbonate, disulfide bonds were reduced in 10 mM dithiothreitol for 30 min, alkylated in the presence of 20 mM iodoacetamide for 30 min in the dark, and digested with trypsin (sequencing grade, Promega) at 1:100 (enzyme: protein ratio) at 37°C overnight while shaking at 1,400 rpm. Beads were dissolved in 95% ACN and shaken at 1,400 rpm for 10 min at RT to bind tryptic peptides to the beads. On the magnetic rack, the supernatant was removed, and beads were washed two times with 100% ACN. Elution was performed with 2% DMSO in 1% formic acid. The supernatant was dried in a vacuum centrifuge and stored at −20 °C until further use.

### Peptide library generation with high pH fractionation

Three human and three mouse brain homogenates were used for library generation. Protein extraction was performed with 7 M urea, 2 M Thiourea, 4% CHAPS, 150 mM DTT, and 0.5% Ampholyte. Lysates were concentrated to a final concentration of 4 mg/mL. For each sample, 50 µg of protein was subjected to tryptic digestion, following the SP3 protocol (Hughes *et al*, 2019). Digests from different samples were combined before fractionation. In total, 50 µg of tryptic peptides were used for High pH RP-HPLC using a 25 cm ProSwift™ RP-4H capillary monolithic column (Thermo Scientific) on an Agilent 1200 series HPLC (high-pressure liquid chromatography) system. A gradient was applied for a total of 45 min with a flow rate of 0.2 mL/min starting at 96.7% eluent A (10 mM NH_4_HCO_3_) and 3.3% eluent B (10 mM NH_4_HCO_3_ in 90% acetonitrile) for 5 min, rising to 38.5% B in 20 min and increasing to 95.0% in 1 min for 10 min and re-equilibrated to 3.3% B for 8 min. Thirty fractions were collected on an Äkta Prime Plus fraction collector, pooled into 13 peptide library fractions, and dried in the vacuum centrifuge.

### Liquid-chromatography-coupled tandem mass spectrometry (LC-MS/MS)

Prior to LC-MS/MS analysis, EV samples and peptide library fractions were dissolved in 0.1% FA to a final concentration of 1 µg/µL. For LC-MS/MS measurements, 1 µg tryptic peptides were injected. Measurements of EV samples were performed on an orbitrap MS (QExactive, Thermo Fisher) coupled to a nano-UPLC (Dionex Ultimate 3000 ultra-performance liquid chromatography system, Thermo Fisher Scientific). Measurements of peptide library fractions were performed on a quadrupole-ion-trap-orbitrap MS (Orbitrap Fusion, Thermo Fisher Scientific) in orbitrap-orbitrap configuration. Chromatographic separation of peptides was achieved with a two-buffer system (buffer A: 0.1% FA in ultrapure H_2_O, buffer B: 0.1% FA in ACN). Attached to the UPLC was a peptide trap (100 μm × 200 mm, 100 Å pore size, 5 μm particle size, C18, Thermo Fisher Scientific) for online desalting and purification followed by a 25 cm C18 reversed-phase column (75 μm × 250 mm, 130 Å pore size, 1.7 μm particle size, Peptide BEH C18, Waters). Peptides were separated using an 80-min gradient with linearly increasing ACN concentration from 2% to 30% ACN in 65 minutes. Eluting peptides were ionized using a nano-electrospray ionization source (nano-ESI) with a spray voltage of 1800 V, transferred into the MS, and analyzed in data-dependent acquisition (DDA) mode. For each MS1 scan, ions were accumulated for a maximum of 240 ms or until a charge density of 1×10^6^ ions (AGC Target) was reached. Fourier-transformation-based mass analysis of the data from the orbitrap mass analyzer was performed, covering a mass range of 400-1,200 m/z with a resolution of 70,000 (at m/z = 200) for EV samples on the orbitrap MS and a resolution of 60,000 for peptide library on the quadrupole-ion-trap-orbitrap MS in orbitrap-orbitrap configuration. Peptides with charge states between 2+ and 5+ above an intensity threshold of 1×10^5^ were isolated within a 2 m/z isolation window from each precursor scan and fragmented with a normalized collision energy of 25% using higher energy collisional dissociation (HCD). MS2 scanning was performed at a resolution of 15,000 for EV samples on the orbitrap MS and a resolution of 17,500 for peptide library on the quadrupole-ion-trap-orbitrap MS in orbitrap-orbitrap configuration, covering a mass range from 100 m/z and accumulated for 50 ms or to an AGC target of 1×10^5^. Already fragmented peptides were excluded for 15 seconds.

### Raw data processing

LC-MS/MS from DDA were searched with the Sequest algorithm integrated into the Proteome Discoverer software (Version 2.4.1.15, Thermo Fisher Scientific) against a reviewed mouse database obtained in October 2020, containing 17,053 entries and a reviewed human database, obtained in June 2021, containing 20,386 entries. Carbamidomethylation was set as a fixed modification for cysteine residues, and the oxidation of methionine and pyro-glutamate formation at glutamine residues at the peptide N-terminus, as well as acetylation of the protein N-terminus, were allowed as variable modifications. A maximum number of two missing tryptic cleavages was set. Peptides between 6 and 144 amino acids were considered. A strict cutoff (FDR<0.01) was set for peptide and protein identification. LC-MS/MS from peptide library fractions were handled in a separate processing step within the software. A multi-consensus workflow was applied to sample and library files (.mgf) to generate a combined output and increase the protein identification rate for individual samples through the feature mapper. Normalization was performed in specific protein amount mode based on the EV marker proteins FLOT1, FLOT2, CD9, CD63, CD37, CD81, CD82, CD151, PDCD6IP, TSG101, RAB1A, RAB1B, RAB2A, RAB2B, RAB3A, RAB3B, RAB4A, RAB4B, RAB5A, RAB5B, RAB6A, RAB6B, RAB7A, RAB7B, RAB8A, RAB8B, RAB9A, RAB9B, RAB10, RAB11B, ANXA1, ANXA2, ANXA3, ANXA4, ANXA5, ANXA6, ANXA7, ANXA11. Protein abundances for individual samples were exported and submitted to subsequent statistical analysis. Protein abundances for library fractions were discarded. Data is available via ProteomeXchange with the identifier PXD045737.

### mRNA content analysis with nCounter^®^ panels (Nanostring)

Samples were processed in the panel without a previous RNA isolation, as published previously (Bub *et al*, 2022). Briefly, BDEVs were isolated from human brain tissues, as explained before (with and without collagenase), and fractions F1 and F2 of the same sample were pooled. PBS was added to the pooled samples and pelleted again for 2 h at 120,000*xg*. After discarding the PBS completely, the BDEVs were resuspended in 10 µL of RTL Lysis Buffer (Qiagen) diluted 1:3 in RNAse-free H_2_O. 5 µL were loaded from each sample in the NanoString nCounter^®^ Neuropathology panel (#XT-CSO-HNROP1-12, NanoString Technologies).

### Statistical and bioinformatics analyses

Statistical analysis was performed with GraphPad PRISM 8 (GraphPad Software, USA) or R. Unless otherwise stated, data are plotted as the mean ± SEM. The normality of the distributions in the NTA-derived results was checked using the Shapiro-Wilk test. If all the samples passed the normality test, the Ordinary one-way ANOVA test and Bonferroni’s multiple comparisons test were performed to validate the statistical differences; if not, the non-parametrical Kruskal-Wallis test was used.

For the statistical and bioinformatics analyses of the proteomics data sets, we mainly relied on Perseus software (Max Planck Institute of Biochemistry, Martinsried, Germany) and various packages in R. Proteins which had at least two valid values in at least one of the groups were kept for further analysis. Initially, proteins with missing values were excluded from the Principal component analyses and heatmaps. Each sample was then normalized by the median protein area per sample, and paired comparisons using Student’s *t*-test were performed using Perseus. Proteins were considered to be significantly different in abundance if the Student’s *t*-test-based *p*-value was ≤ 0.05 and at least a 1.5-fold change in either direction was observed. Proteins were considered uniquely present in a certain group when they were expressed in at least two out of three replicates for that particular group and were simultaneously absent in the other group during pairwise comparison.

Principal component analysis (PCA) was carried out using the prcomp function of the base R. Resultant PCA plots were composed utilizing the ggplot2 (version 3.3.3; Wickham, H. 2016). Hierarchical clustering and heatmaps were prepared using heatmp.2 function in the gplot package (version 3.1.1. Warnes, G.R. et al, 2021). Correlation plots were prepared using ggcorrplot package (version 0.1.4.; Kassambara, A. 2021). Peason’s coefficient values were obtained using base R. Gene ontology (GO) analyses were performed utilizing the clusterProfiler, enrichR, org.Hs.eg.db, org.Mm.eg.db, and topGO packages. Overrepresentation analysis (ORA) was performed to find associations of differentially upregulated proteins along with proteins uniquely detected in F1 (+ and −) and F2 (+ and −) individually with the GO category ‘Cellular Components’ with the following analysis parameters: *p*-valueCutoff = 0.05, *q*-valueCutoff = 0.2, minGSSize = 5, and maxGSSize = 500.

Proteins constituting the EV preparations were checked for their presence in the Vesiclepedia database (Kalra *et al*, 2012; Pathan *et al*, 2019), using FunRich enrichment software (Pathan *et al*, 2015) on 10^th^ December 2023. Mouse proteins were compared to the human Vesiclepedia database, as the corresponding mouse database was not supported on the FunRich platform.

For the nCounter^®^ panel analysis, all initial statistical work-up was performed using nSolver analysis software (version 4.0.70). For the downstream analyses of mRNA expression profiles, genes with raw expression scores not exceeding the average plus two standard deviations of all corresponding negative control probes were removed from the analysis. All expression values below the 20 reads threshold were imputed to 20. Normalization was based on the tetraspanin mRNAs present on the panel i.e. *CD14, CD33, CD34, CD4, CD40, CD44, CD68, CD8A,* and *CD9.* All genes with no detection in any of the samples were excluded from the analysis. An mRNA was considered differentially expressed if the corresponding absolute log2-fold change (log2FC) was ≥ 0.584 and the *p*-value ≤ 0.05. PCA, heatmaps, and correlation plots were prepared as those for proteomics, as described above.

## Results

### Collagenase treatment of brain homogenates impact on key BDEV proteins, including PrP^C^

In a previous study, our groups reported the enrichment of the proteolytically truncated PrP^C^-C1 fragment (generated by physiological α-cleavage (Linsenmeier *et al*, 2017)) in BDEVs and highlighted the proteolytic impact of the enzymatic tissue digestion (employed during BDEV isolation) on the PrP^C^-C1/flPrP^C^ ratio (Brenna *et al*, 2020). In order to further examine this effect we first digested mouse brain tissue with collagenase D and collagenase III, (both widely used in EV isolation protocols from tissue e.g. (Su *et al*, 2021; Crescitelli *et al*, 2021)) (Figure 1a). Tissue was lysed with PBS, and both low and high doses of collagenase D and III (high dose = 40 U/mL, low dose = 10 U/mL) were used. As a control, we included non-digested homogenate with and without the addition of protease inhibitors to exclude endogenous protease activity. The results indicated that the enzymes cleaved the EV-related proteins Alix, CD9, and CD81, while 14-3-3, ADAM10, LRP1, and Flotillin-1 remained unaffected (Figure 1b). Interestingly, GM130 clearly disappeared in all the enzymatic treatments, highlighting its sensitivity to collagenase digestion. We also conducted the same experiment using human brain tissue, which also demonstrated protein pattern modifications in CD81 and Alix but not CD9, suggesting different effects of the collagenases depending on the protein sequence variability among species (Supplementary Figure S1a). We then specifically examined the effect of collagenase on PrP^C^ cleavage due to the relevance of the different PrP^C^ forms in its physiological functions (reviewed in (Linsenmeier *et al*, 2017; Mohammadi *et al*, 2023). After digestion, the homogenate was treated with PNGase to remove the glycans and enable more clear distinctions in PrP^C^ patterns. Western blotting analysis using two anti-PrP^C^ antibodies revealed two cleavage sites: one comparable to the physiological α-cleavage and another artificially generated by collagenase in the N-terminal region (Figure 1c). The same protein digestion pattern was observed when human brain homogenate was tested (Supplementary Figure S1b). Uncropped blots and corresponding total protein staining are included in Supplementary Figure S2a-b.

### Unlocking collagenase’s impact on EVs: membrane EV proteins are Cleaved, whereas intraluminal ones are spared

To discard the possibility of artifact generation resulting from EVs isolation from a complex system such as brain tissue, we examined the effect of collagenase D on isolated EVs from cultured N2a cells (N2a-EVs). The N2a-EVs were isolated by filtration and ultracentrifugation (Figure 1d). NTA analysis showed the usual size distribution with a concentration of 9.17×10^11^ ± 1.54×10^11^ particles per mL with a mode size of 139,6 ± 2.13 nm (Figure 1e-f). The typical cup shape of N2a-EVs was confirmed using TEM (Figure 1g).

N2a-EVs (3×10^10^) were incubated with collagenase D (2 mg/mL, equivalent to 40 U/mL) for 30 min, and the resulting samples were subjected to western blot analysis. CD81 and PrP^C^ were found to be clearly cleaved, generating “artificial” protein fragments, whereas the EV-luminal proteins Alix and Flotillin-1 remained much less affected. These proteins, along with ADAM10, just showed slightly lower levels due to either the disruption of EVs during isolation and the collagenase digestion or the digestion continuing after the EV lysis for western blotting. In line with this, we have previously observed that even a high concentration of protease inhibitors cannot effectively prevent collagenase activity (unpublished data). The Golgi protein GM130 was absent in all N2A-EV samples (Figure 1h). For uncropped blots and corresponding total protein staining, see Supplementary Figure S2c.

These results highlight the effect of collagenase on unspecific trimming and degradation of EV-associated and ND-related proteins, which could drastically affect downstream analyses and functionality of isolated BDEVs.

### Characterization of a simplified and non-enzymatic BDEV isolation protocol

In order to mitigate the protein cleavage effect of collagenase digestion, we tested the possibility of isolating BDEVs with a reduced amount of collagenase D (0.5 mg/mL) and without collagenase. Moreover, to reduce the protease activity, we also reduced the enzymatic digestion time and included the addition of PI. We performed the BDEV isolation by slightly modifying a previously published protocol to isolate EVs from tumor tissues (Crescitelli *et al*, 2021) as schematically shown in Figure 2.

**Figure 2:**
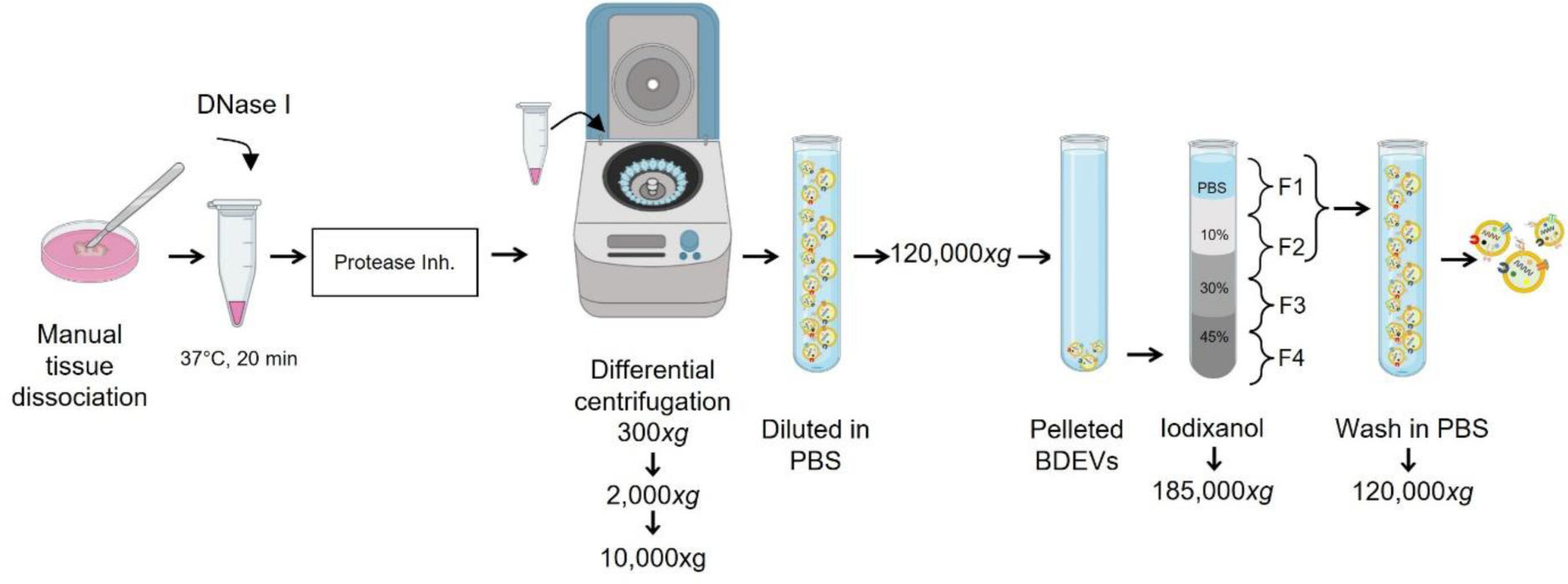
Simplified non-enzymatic BDEV isolation protocol. Schematic representation of the new enzyme-free BDEVs isolation protocol: the brain tissue was manually minced in RPMI media and transferred to a new tube where a DNAse I treatment was carried out at 37°C for 30 min. The reaction was stopped by adding a protease inhibitor cocktail. Then differential centrifugation was applied (200*xg*, 2,000*xg*, and 10,000*xg*), and the BDEV-containing supernatant was diluted in PBS. The BDEVs were pelleted and washed at 120,000*xg* before being separated in an iodixanol-based gradient at 185,000*xg*. Fractions were collected, and the F1 and F2 (which mainly contained the BDEVs) were washed in PBS and pelleted at 120,000*xg*. Figure created with BioRender.com.

To evaluate the efficiency of our protocol in different systems, we isolated EVs from mouse and human brain tissues. Each brain tissue sample was cut into two pieces (100-180 mg per piece): one piece was isolated without the addition of collagenase (w/o), while the other underwent isolation with collagenase (0.5 mg/mL). The BDEV characterization yielded consistent results in both mouse and human samples (Figures 3 and 4). A clear expression of the positive EV markers (EV+) Alix, CD81, and Flotillin-1 was observed with both protocols. However, CD81 and Flotillin-1, as observed in Figure 1, exhibited distinct patterns in the presence of collagenase. Moreover, the EV-negative marker Lamin A/C was not detected in any fraction, but GM130 was present in F2 of the BDEVs isolated without collagenase, suggesting that its absence in the collagenase-treated samples is due to digestion (as observed in Figure 1 for tissue homogenates). Notably, PrP^C^ was predominantly found in F2 of mouse-derived BDEVs but distributed across both fractions of the human BDEVs. Regardless of the source, PrP^C^ exhibited cleaved patterns in the enzyme-based BDEVs samples, indicating an enriched PrP-C1-like fragment (Figures 4a and 5a) consistent with our observations in the brain homogenate digestion (Figure 1c). To determine whether the proteolytic changes observed were directly linked to the enzymatic isolation, we checked protein levels in the 10K pellet and the Pre-gradient BDEVs (Supplementary Figures S3a and S4a). The PrP^C^ pattern was similar to the one observed in F1 and F2, with an enriched PrP-C1-like fraction in the collagenase-based BDEVs. GM130 was observed in the enzyme-free BDEVs samples, but not in the enzyme-based ones, again suggesting that the absence of GM130 in these samples is due to digestion (Supplementary Figures S3a and S4a). As expected, no EV markers were detectable in the F3 and F4 fractions, and only a weak GM130 signal in the mouse BDEV was observed (Supplementary Figures S3b and S4b). Uncropped blots and corresponding total protein staining are presented in Supplementary Figure S5.

**Figure 3:**
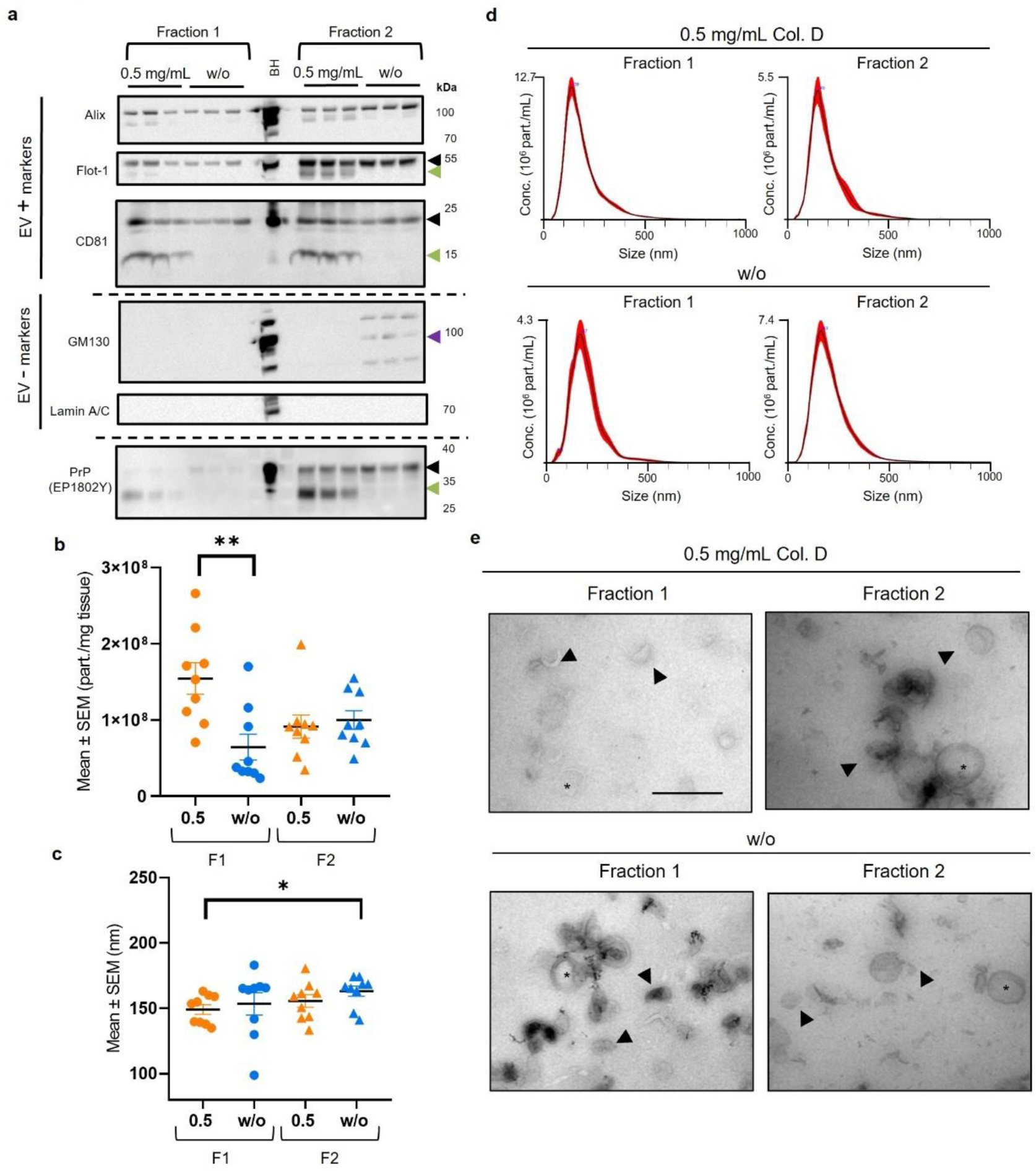
BDEV isolation from mouse brain tissue without enzymatic digestion maintains BDEV purity and prevents artificial proteolytic processes. (a) Western blots of F1 and F2 BDEVs samples isolated from mouse brain tissue, with or without the addition of 0.5 mg/mL of collagenase D (*n* = 3 for each condition), were labeled for PrP^C^, the EV positive markers Alix, Flotillin-1, and CD81 as well as for the Golgi and nucleus markers, GM130 and Lamin A/C as EV negative markers. The green arrowheads indicate the artificial cleavage observed in the proteins PrP^C^, CD81, and Flotillin-1, while the purple arrowhead denotes the unexpected presence of GM130. A total mouse brain homogenate (BH) served as a control. The concentration of particles per mg of initial tissue (b) and mode size analysis in nm (c) of BDEVs obtained with both protocols were measured with NTA (*n* = 9 for each condition). No differences in the particle mode size were observed between the same fraction in both protocols (just the F1^+^ has smaller EVs than F2^−^), but the F1^−^ displayed significantly fewer particles/mg of tissue than the F1^+^. (d) Representative size distribution graphs from NTA analysis of F1 and F2 BDEVs show the expected normal-like distribution. (e) TEM images of negative stained BDEV showing the typical double membrane and cup shape. Small BDEVs are indicated with arrowheads, while larger vesicles are marked with asterisks (*). Scale bar = 500 nm. Data are presented as mean ± S.E.M. in (b) and (c).

**Figure 4:**
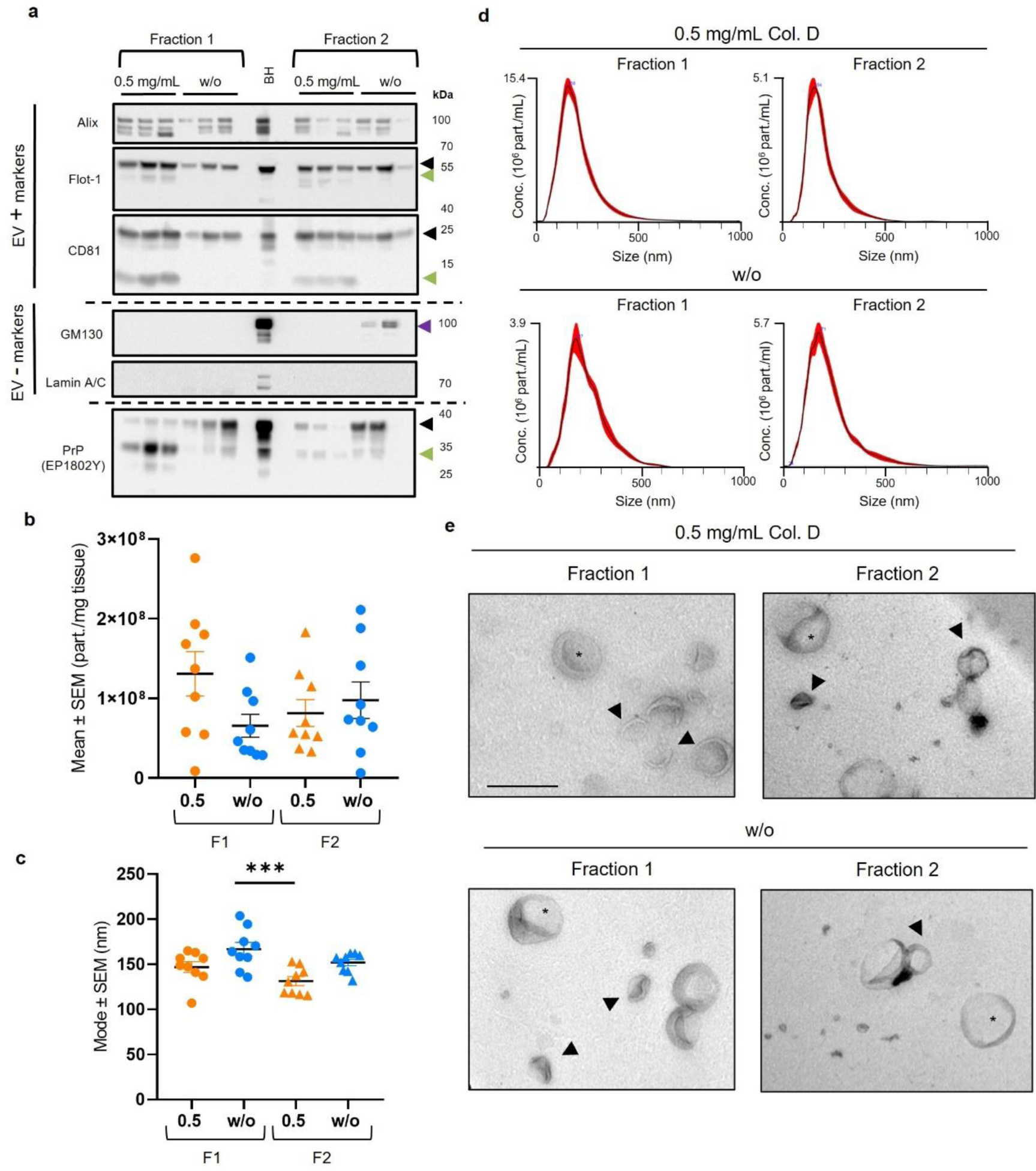
BDEV isolation from human brain tissue without enzymatic digestion maintains BDEV purity and prevents artificial proteolytic processing. (a) Representative western blots of F1 and F2 BDEVs samples isolated from human brain tissue, with or without the addition of 0.5 mg/mL of collagenase D (*n* = 3 for each condition), for PrP^C^, the EV positive markers Alix, Flotillin-1, and CD81 and the Golgi and nucleus markers, GM130 and Lamin A/C, as EV negative markers. The green arrowheads indicate the artificial cleavage observed in the proteins PrP^C^, CD81, and Flotillin-1, and the “unexpected” bands for GM130 (indicated by purple arrowhead). A total human brain homogenate (BH) was used as a loading control. The concentration of particles per mg of initial tissue (b) and mode size analysis in nm (c) of BDEVs obtained with both protocols were measured with NTA (*n* = 9 for each condition). No differences were observed in the particle concentration between both protocols, however, the F2^+^ has a significantly larger particle size than the F1^−^. (d) Representative size distribution graphs from NTA analysis of F1 and F2 BDEVs show the expected normal-like distribution. (e) TEM images of negative stained BDEV illustrating the typical double membrane and cup shape. Small BDEVs are indicated with arrowheads, and larger vesicles are marked by the asterisks (*). Scale bar = 500 nm. Data are presented as mean ± S.E.M. in (b) and (c).

**Figure 5:**
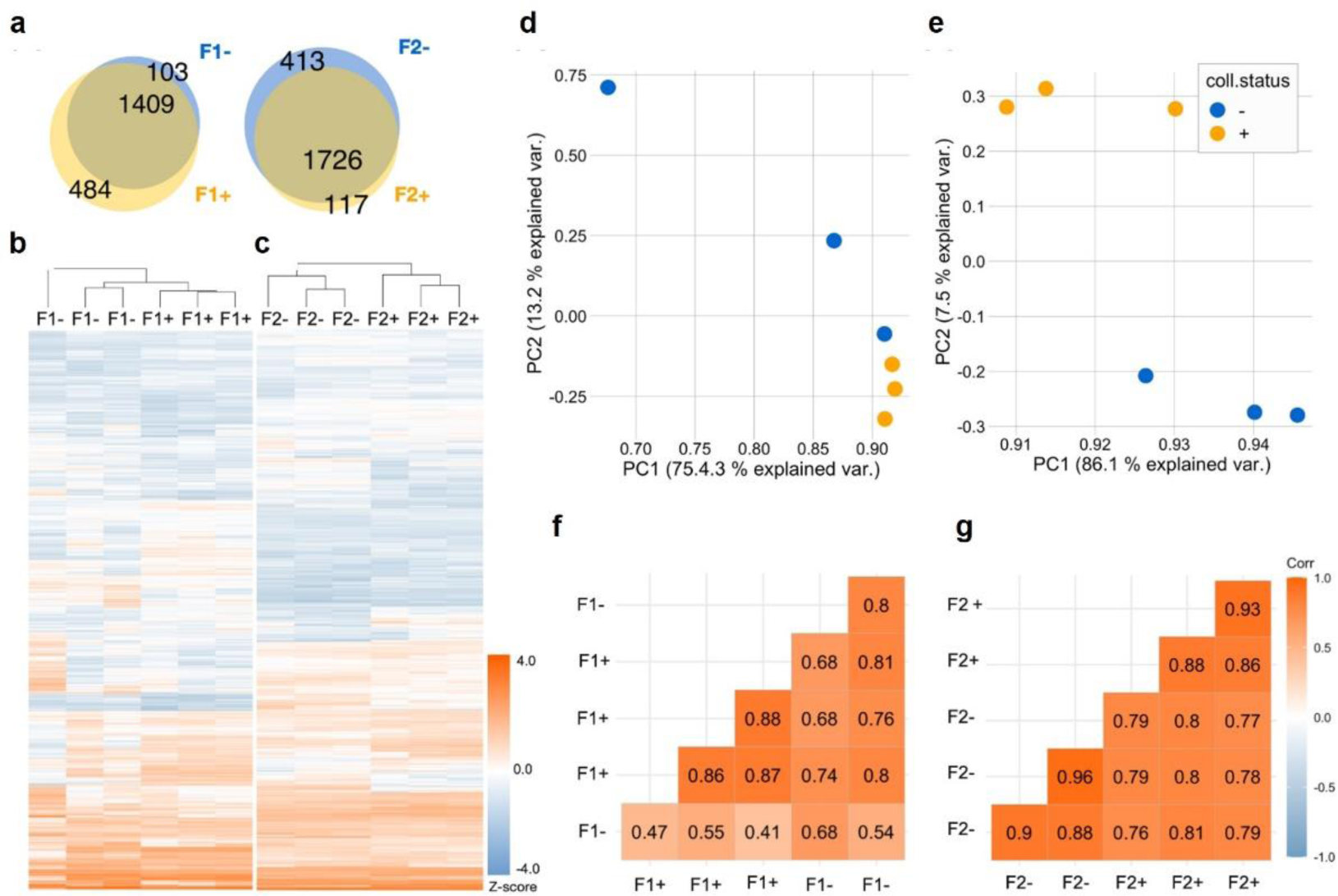
Systematic analysis of mouse BDEV proteomes: no major differences are observed between BDEVs isolated enzyme-free or with collagenase D. The proteomes of mouse BDEVs isolated using the collagenase-free method (indicated with ‘-’) and those with the collagenase-based protocol (labeled as ‘+’) were analyzed systematically (n = 3 for each condition). (a) Venn diagrams display a high compositional overlap between F1^−^ and F1^+^ as well as between F2^−^ and F2^+^. 1,512 proteins were found in F1^−^, 1,893 in F1^+^, 2,139 in F2^−^ and 1,843 in F2^+^. A total number of 1409 proteins were found common between F1^−^ and F1^+^, where 103 proteins were uniquely present in F1^−^and 484 proteins were uniquely found in F1^+^. In the case of F2, 1,726 were found to be common, 413 proteins were found uniquely in F2^−^ and 117 proteins were only present in F2^+^. Heatmaps displaying protein abundances (column z-score) of F1^−^ compared to F1^+^ (b) and F2^−^ versus F2^+^ (c). Scatter plots showing principal component analysis (PCA) of the proteomic composition of mouse BDEVs prepared using the two protocols for F1 (d) and F2 (e). Correlation plots highlighting the similarities between F1 (-vs +) (f) and F2 (-vs +) (g). The color key represents variations in Pearson’s correlation coefficient values.

NTA revealed particle counts between 0.3-2.8×10^8^ particles/mg of tissue in both mouse and human isolates, indicating a comparable BDEV isolation yield (Figures 3b and 4b). In the case of mouse BDEV, there were significantly more particles/mg in the F1 of collagenase samples (F1^+^) compared to the F1 enzyme-free samples (F1^−^), while all other fractions had a very similar number of particles (F1^+^= 1.54×10^8^ ± 2.06×10^7^ part./mg; F2^+^= 9.1×10^7^ ± 1.52×10^7^ part./mg; F1^−^= 6.4×10^7^ ± 1.68×10^7^ part./mg; and F2^−^= 9.9×10^7^ ± 1.21×10^7^ part./mg - F1^+^vs F2^+^: *p* = 0.201; F1^+^vs F1^−^: *\*\*p* = 0.004; F1^+^vs F2^−^: *p* = 0.46; F1^−^vs F2^+^: *p* > 0.99; F1^−^vs F2^−^: *p* = 0.61; F2^+^vs F2^−^: *p* > 0.99, Kruskal-Wallis test) (Figure 3b). The NTA analysis also revealed no difference when comparing the same fractions in terms of the size of the particles, exhibiting the usual size distribution (Figure 3d). The only significant difference in particle size was detected between F1^+^ and F2^−^, which is expected since the EVs are collected from different density fractions (F1^+^= 149.1 ± 3.7 nm; F2^+^= 155.6 ± 4.8 nm; F1^−^= 153.4 ± 8.5 nm; F2^−^= 163.1 ± 3.9 nm – F1^+^vs F2^+^: *p* > 0.99; F1^+^vs F1^−^: *p* = 0.70; F1^+^vs F2^−^: \**p* = 0.043; F1^−^vs F2^+^: *p* > 0.99; F1^−^vs F2^−^: *p* > 0.99; F2^+^vs F2^−^: *p* = 0.76, Kruskal-Wallis test) (Figure 3c).

Likewise, no differences were observed in terms of particle concentration of the human BDEVs in any fraction (F1^+^= 1.31×10^8^ ± 2.78×10^7^ part./mg; F2^+^= 8.15×10^7^ ± 1.68×10^7^ part./mg; F1^−^= 6.55×10^7^ ± 1.45×10^7^ part./mg; and F2^−^= 9.77×10^7^ ± 2.30×10^7^ part./mg - F1^+^vs F2^+^: *p* = 0.66; F1^+^vs F1^−^: *p* = 0.22; F1^+^vs F2^−^: *p* > 0.99; F1^−^vs F2^+^: *p* > 0.99; F1^−^vs F2^−^: *p* > 0.99; F2^+^vs F2^−^: *p* > 0.99, Bonferroni’s multiple comparison tests, Ordinary one-way ANOVA) (Figure 4b). Furthermore, NTA revealed the usual size distribution of the human BDEV (Figure 4d), with a mode size just significantly different between the F2^+^ and the F1^−^, which, is likely since they come from different density fractions (F1^+^= 148.6 ± 5.9 nm; F2^+^= 131.4 ± 5.0 nm; F1^−^= 166.8 ± 7.4 nm; F2^−^= 152.0 ± 3.5 nm - F1^+^vs F2^+^: *p* = 0.37; F1^+^vs F1^−^: *p* = 0.11; F1^+^vs F2^−^: *p* > 0.99; F1^−^vs F2^+^: *\*\*\*p* = 0.0006; F1^−^vs F2^−^: *p* = 0.44; F2^+^vs F2^−^: *p* = 0.09, Bonferroni’s multiple comparison tests, Ordinary one-way ANOVA) (Figure 4c).

Overall, the NTA results demonstrated that using the collagenase-free protocol yields particles with size distribution and concentration of particles per tissue weight similar to the samples isolated with 0.5 mg/mL of collagenase D (Figure 4d).

Lastly, we characterized BDEVs by labeling them with negative staining (uranyl acetate) and imaged them with TEM (Figures 3e and 4e). We observed the characteristic cup-shaped and membranous EV structures in both fractions (F1 and F2), with no discernible differences between the protocols or tissue origins. For a comprehensive illustration of the variety of BDEVs obtained across fractions and protocols, please refer to Supplementary Figure S6, which presents a detailed compilation of TEM images.

In order to assess if our protocols (with reduced collagenase or without any protease added) obtained a different concentration of BDEV due to their reduced enzyme treatment, we compared BDEVs (from mouse tissue) isolated with the reference protocol described by Crescitelli *et al* with 2 mg/mL of collagenase D (with some minor modifications to adapt it to our equipment) (Crescitelli *et al*, 2021). The isolated BDEVs expressed the EV markers Alix, CD81, and Flotillin-1 but not the nuclear and Golgi markers Lamin A/C and GM130, respectively. However, as described in Figure 3, CD81, Flotillin-1, and PrP^C^ showed clear cleaved protein patterns again (Supplementary Figure S7a). The NTA analysis showed the characteristic size distribution in both fractions (Supplementary Figure S7b), but no differences in concentration or size compared to BDEVs isolated with less (0.5 mg/mL) or without collagenase (Supplementary Figure S7c-d). These results demonstrate that our approaches do not result in a reduced BDEV yield.

### The non-enzymatic BDEV isolation protocol minimally impacts the EV-related proteome but alters some proteins that may be associated with the EV membrane and corona

To assess whether a different population of EVs was being isolated with and without collagenase, we conducted a proteomic analysis of BDEVs from both mouse and human brain tissue samples either isolated with 0.5 mg/mL of collagenase D or without enzymatic treatment. For the analysis, we paid special attention to proteins known to be in the EV membrane and corona. Overall, the major differences were observed between the F1 and F2 fractions in both instances, isolated with the collagenase-free (F1^−^ and F2^−^) and with collagenase-based (F1^+^ and F2^+^) protocols, in both mouse and human BDEVs (Figures 5, 6, 7, and 8).

No major differences were observed for the mouse BDEV (both F1 and F2) prepared with or without collagenase, displaying high inter-group similarities in the proteomic profiles, as depicted by protein abundance heatmaps and associated hierarchical clustering (Figures 5b and 5c). Out of 1,512 proteins found in F1^−^ and 1,893 found in F1^+^, 1,409 were common in both F1 samples, 103 were just found on the F1^−^, and 484 were unique for F1^+^. In the case of F2, from 2,139 proteins found on F2^−^ and 1,843 in F2^+^, 1,726 were commonly found, 413 were just found in F2^−^, and 117 in F2^+^ (Figure 5a), thus following a similar pattern as F1.

PCA plots of commonly expressed proteins for F1^−^ and F1^+^ (principal component (PC)-1 and PC-2 representing 75.4% and 13.2% of data variances, respectively) and correlation plots with high Pearson’s coefficient values highlight a high degree of overlap between their proteomic compositions (Figures 5d and 5f). However, one F1^−^ sample behaved as an outlier, as depicted by lower Pearson coefficient values (Figure 5f). Related to F2^−^ and F2^+^ (with PC-1 and PC-2 representing 86.1% and 7.5% of data variances, respectively), the correlation plot also displayed resemblances in the proteomic profiles (Figures 5e and 5g).

When checked for bona fide EV marker proteins (defined by literature) between both F1 and F2 preparations no substantial changes between collagenase-free and collagenase-based methods were observed, showing that the BDEV populations isolated were similar (Figures 6a and 6e). In pairwise comparisons, 84 proteins were found to be relatively enriched in F1^−^, while 94 proteins displayed lower expression in the F1^−^ samples (Figure 6b). The proteins with increased abundance in the F1^−^ were strongly associated with the following GO terms (cell components): neuron to neuron synapse (GO:0098984), postsynaptic density (GO:0014069), asymmetric synapse (GO:0032279), postsynaptic specialization (GO:0099572), cytoplasmic region (GO:0099568), synaptic vesicle membrane (GO:0030672), exocytic vesicle membrane (GO:0099501), synaptic vesicle (GO:0008021), endoplasmic reticulum tubular network membrane (GO:0098826), and transport vesicle membrane (GO:0030658) (Figure 6c); whereas proteins comparatively less expressed in the F1^−^, were found associated with synaptic vesicle membrane (GO:0030672), exocytic vesicle membrane (GO:0099501), coated vesicle (GO:0030135), proton-transporting V-type ATPase complex (GO:0033176), transport vesicle membrane (GO:0030658), synaptic vesicle (GO:0008021), clathrin-coated vesicle membrane (GO:0030665), transport vesicle (GO:0030133), exocytic vesicle (GO:0070382), and vacuolar proton-transporting V-type ATPase complex (GO:0016471) (Figure 6d).

**Figure 6:**
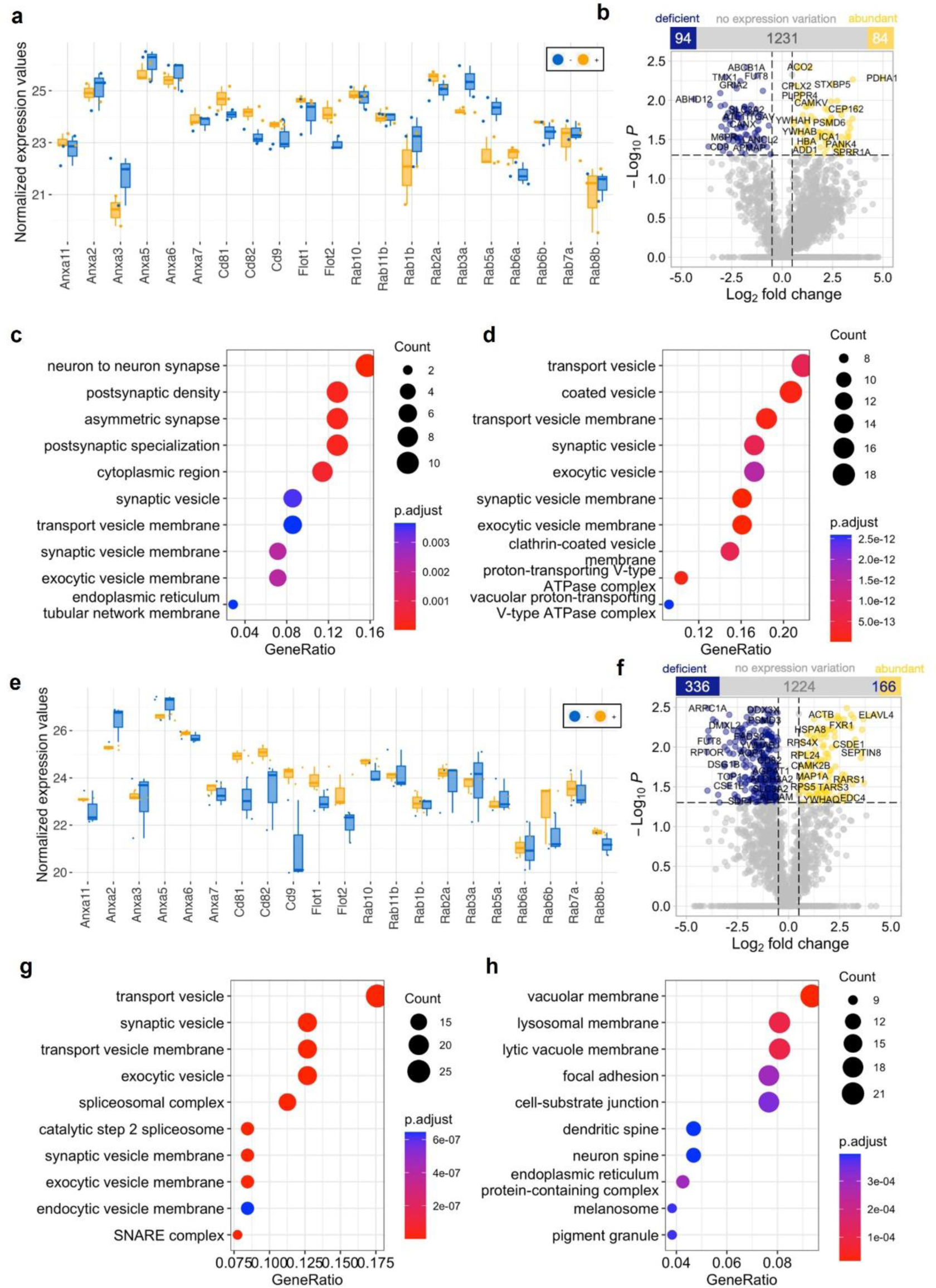
No significant differences in bona fide EV marker proteins were observed when comparing the proteomic profiling of BDEVs isolated from mouse brain tissue in density fractions 1 and 2 prepared with or without collagenase. (a) Plot showing normalized expression values of 21 bona fide EV marker proteins. No significant differences were found between the col+ or col-samples. (b) Volcano plot showing proteins relatively down (shown in blue) and up-regulated (shown in yellow) in F1^−^ compared to F1^+^. 84 proteins were found to be overexpressed, and 94 proteins were found to be downregulated in F1^−^ compared to F1^+^, as shown in the column graph associated with the volcano plot. The intercepts on the x-axis indicate the cut-offs for expression changes, set at −1.5 fold for down-regulation and +1.5 fold for up-regulation. The intercept on the y-axis marks the cut-off set for a *p*-value of 0.05. (c and d) Dot plots depicting the top 10 cellular component categories linked to proteins up (c) and downregulated (d) in F1^−^. These categories are based on gene ontology terms, and the color key indicates the adjusted *p*-values resulting from the over-representation analysis. (e) Normalized expression values of the bona fide EV marker proteins. (f) Volcano plot showing proteins relatively under-represented (shown in blue) and up-regulated protein (shown in yellow) in F2^−^. 166 proteins were found to be more abundant, and 336 proteins were found to be less expressed in F2^−^ when compared to F2^+^. The intercepts along the x-axis represent the thresholds for expression changes: −1.5 fold for down-regulation and +1.5 fold for up-regulation. Meanwhile, the intercept on the y-axis indicates the threshold set for a p-value of 0.05. (g and h) Dot plots depicting the top 10 cellular component categories linked to proteins up (g) and downregulated (h) in F2^−^. These categories are based on gene ontology terms, and the color key indicates the adjusted *p*-values resulting from the over-representation analysis.

In the context of F2^−^ versus F2^+^ comparison, 166 proteins were identified to be significantly more abundant in F2^−^, and 336 proteins were found to be significantly under-represented in F2^−^ compared to F2^+^ BDEVs (Figure 6f). GO analysis revealed that the upregulated proteins in the F2^−^ (and F2^−^ unique proteins) were highly associated with transport vesicle (GO:0030133), synaptic vesicle (GO:0008021), SNARE complex (GO:0031201), transport vesicle membrane (GO:0030658), exocytic vesicle (GO:0070382), spliceosomal complex (GO:0005681), catalytic step 2 spliceosome (GO:0071013), synaptic vesicle membrane (GO:0030672), exocytic vesicle membrane (GO:0099501), and endocytic vesicle membrane (GO:0030666); where the down-regulated proteins were majorly associated with vacuolar membrane (GO:0005774), lysosomal membrane (GO:0005765), lytic vacuole membrane (GO:0098852), endoplasmic reticulum protein-containing complex (GO:0140534), focal adhesion (GO:0005925), cell-substrate junction (GO:0030055), melanosome (GO:0042470), pigment granule (GO:0048770), dendritic spine (GO:0043197), and neuron spine (GO:0044309) (Figures 6g and 6h).

In the case of human BDEVs, hierarchical clustering of the proteomic profiles of the BDEV isolated with the two methods and in both F1 and F2, described a close inter-group similarity, as shown in the protein abundance heatmaps (Figures 7b and 7c). Out of 1,899 proteins identified in F1^−^ and 2,204 found in F1^+^, 1,794 were common in both F1 samples, 105 were exclusively found in F1^−^, and 410 were unique for F1^+.^ In the case of F2, similar trends were found: from 1,905 detected proteins in F2^−^ and 2,158 in F2^+^, 1,754 were commonly found, 151 were just in F2^−^, and 404 in F2^+^ (Figure 7a). In the PCA, BDEVs from F1 (F1^−^ and F1^+^) and F2 (F2^−^ and F2^+^) exhibited close clustering (Figures 7d and 7e) along with high Pearson’s correlation coefficient values (Figures 7f and 7g), highlighting that only subtle differences could be observed among the collagenase-free and collagenase-based preparations.

**Figure 7:**
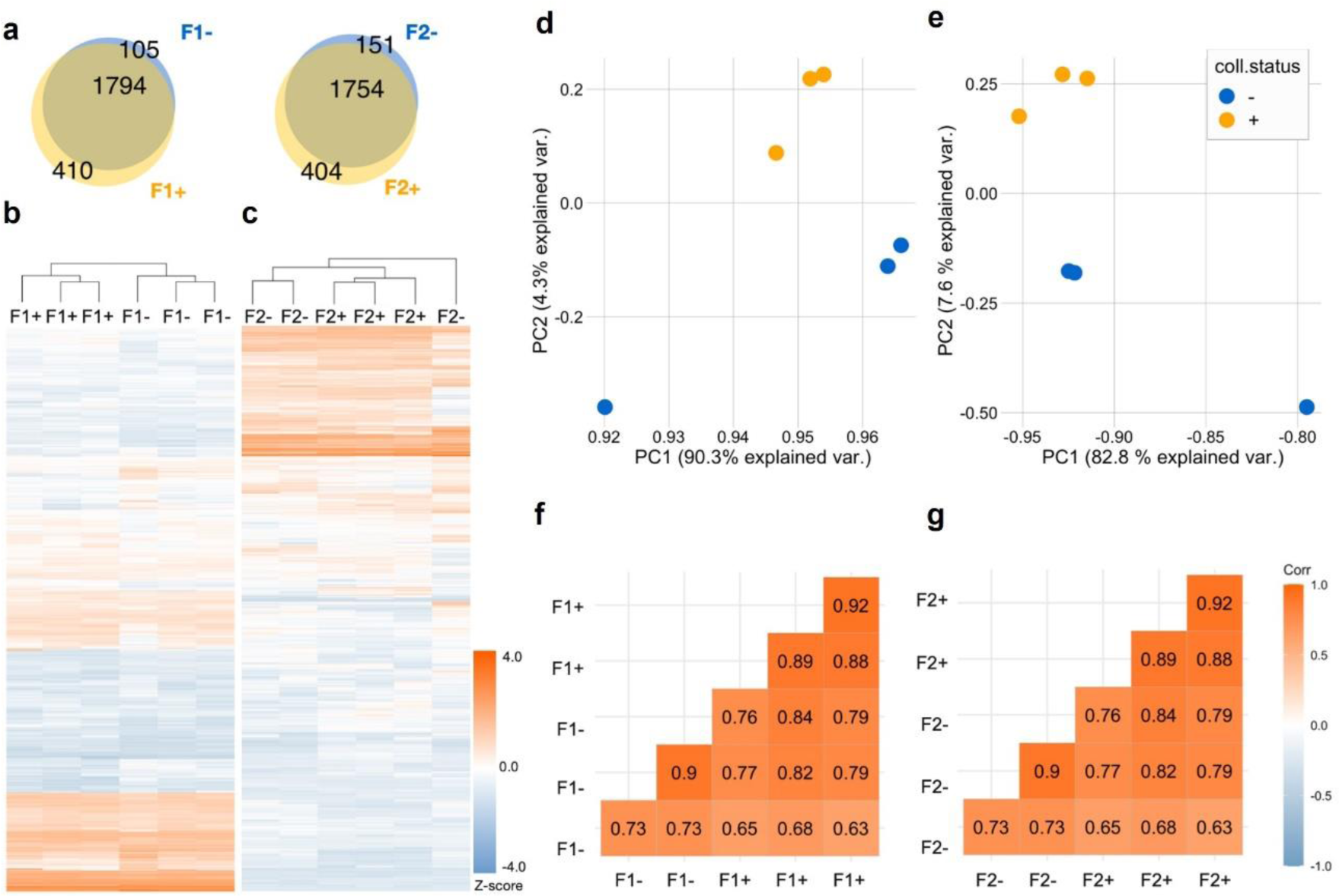
No major differences in the proteomic analysis of BDEVs from human samples comparing isolation with and without collagenase. The proteomes of human-BDEVs isolated using the collagenase-free method (indicated with ‘-’) and those with the collagenase-based protocol (labeled as ‘+’) were analyzed systematically (n = 3 for each condition). (a) Venn diagrams display a high compositional overlap between F1^−^ and F1^+^ as well as between F2^−^ and F2^+^. A total of 1,794 proteins were found common between F1^−^ and F1^+^, where 105 proteins were uniquely present in F1^−^ samples and 410 proteins were uniquely found in F1^+^. In the case of F2, 1754 were found common between F2^−^ and F2^+^. 151 proteins were found uniquely in the F2^−^ group and 404 proteins were only present in the F2^+^. Heatmaps displaying protein abundances (column z-score) of F1^−^ compared to F1^+^ (b) and F2^−^ versus F2^+^ (c). Scatter plots showing principal component analysis (PCA) of the proteomic composition of mouse-derived BDEVs prepared using the two protocols for F1 (d) and F2 (e). The correlation plots highlight the similarities between F1 (-vs +) (f) and F2 (-vs +) samples (g). The color key represents variations in Pearson’s correlation coefficient values.

As observed with the mouse-derived BDEVs, no expression variations for the bona fide EV were found between the collagenase-free and collagenase-based human BDEV in both the F1 and the F2, validating again the isolation of similar BDEV populations (Figures 8a and 8e). We found that there were 191 proteins relatively more abundant in the F1^−^, and 38 were relatively downregulated in F1^−^ compared with the F1^+^ samples (Figure 8b). Related to gene ontology analysis, we found that proteins abundant in the F1^−^ were associated prominently with the postsynaptic specialization (GO:0099572), postsynaptic density (GO:0014069), transport vesicle (GO:0030133), asymmetric synapse (GO:0032279), synaptic vesicle (GO:0008021), cell cortex (GO:0005938), neuron to neuron synapse (GO:0098984), exocytic vesicle (GO:0070382), microtubule (GO:0005874), transport vesicle membrane (GO:0030658) (Figure 8c). Whereas the proteins that were abundantly present in the F1^+^ samples were associated majorly with the inner mitochondrial membrane protein complex (GO:0098800), mitochondrial protein-containing complex (GO:0098798), mitochondrial inner membrane (GO:0005743), respiratory chain complex (GO:0098803), mitochondrial respirasome (GO:0005746), respirasome (GO:0070469), mitochondrial respiratory chain complex I (GO:0005747), NADH dehydrogenase complex (GO:0030964), respiratory chain complex I (GO:0045271), oxidoreductase complex (GO:1990204) (Figure 8d).

**Figure 8:**
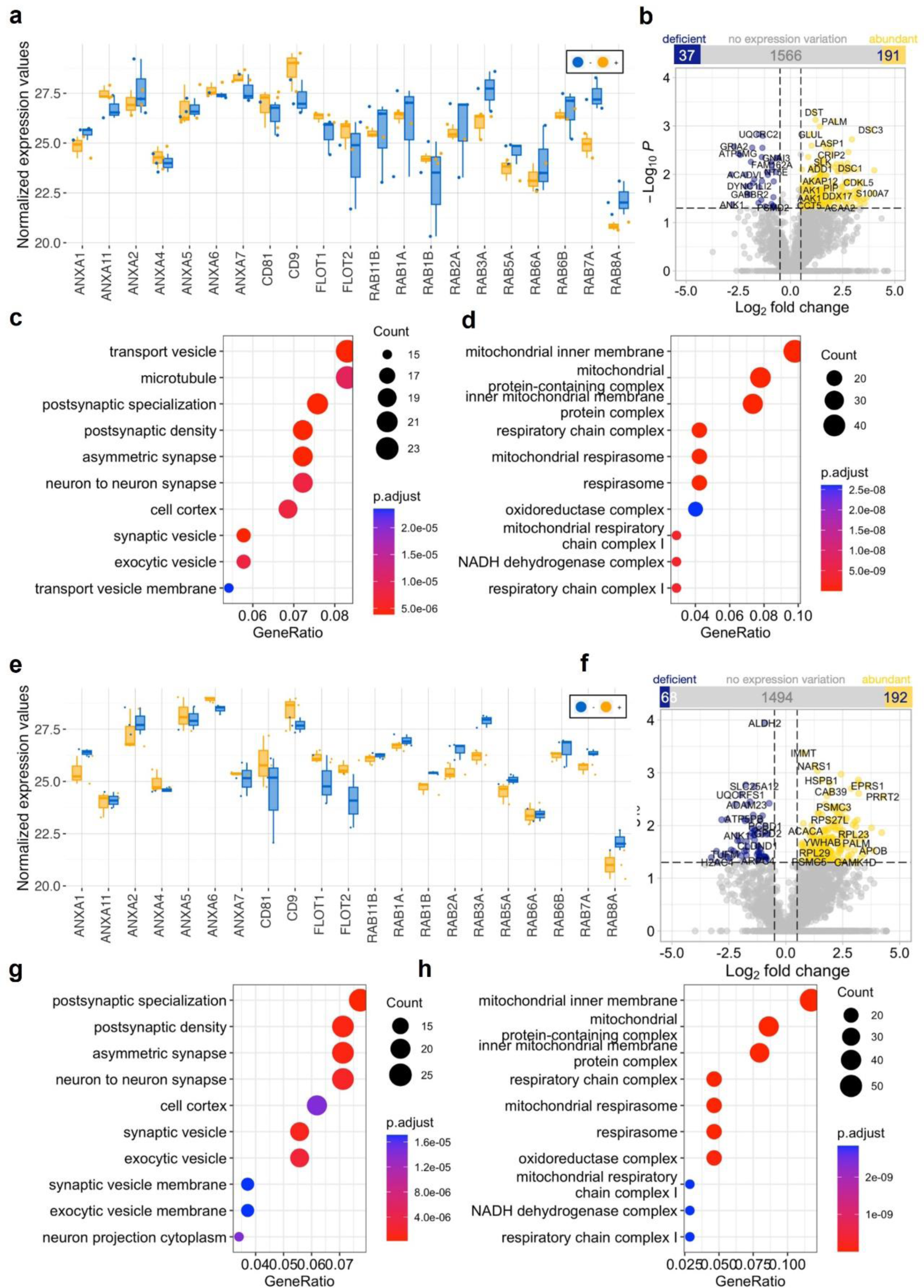
No significant differences in bona fide EV marker proteins were observed when comparing the proteomic profiling of human BDEVs in density fractions 1 and 2 isolated with or without collagenase. Normalized expression values of 21 bona fide EV marker proteins showed no significant differences between col+ or col-samples on F1 (a) or F2 (e) comparisons. (b) Volcano plot showing proteins relatively under-represented (displayed in blue) and up-regulated in F1^−^ (displayed in yellow). 191 proteins were found to be more abundant, and 37 proteins were found to be less expressed in F1^−^, as shown in the column graph associated with the volcano plot. Cut-offs were set at −1.5 fold for down-regulation and +1.5 fold for up-regulation. The intercept on the y-axis marks the cut-off set for a *p*-value of 0.05. (c and d) Dotplots displaying the top 10 categories of cellular components associated with the proteins abundant in F1^−^ abundant and deficient in F1^−^, respectively (gene ontology terms, the color key represents the adjusted *p*-values from over-representation analysis). (f) Volcano plot showing proteins relatively down (displayed in blue) and up-regulated in F2^−^ EVs (displayed in yellow). The intercepts along the x-axis represent the thresholds for expression changes: −1.5 fold for down-regulation and +1.5 fold for up-regulation. Meanwhile, the intercept on the y-axis indicates the threshold set for a *p*-value of 0.05. 192 proteins were found to be increased, and 68 proteins were found to be decreased in F2^−^, shown in the column graph associated with the volcano plot. (g and h) Dotplots displaying the top 10 categories of cellular components associated with the proteins abundant and deficient in F2^−^, respectively (gene ontology terms, color key represents the adjusted *p*-values from over-representation analysis).

Likewise, 192 proteins were significantly upregulated in F2^−^, and 68 proteins were downregulated in F2^−^ compared to F2^+^ BDEVs (Figure 8f). For F2^−^ BDEVs associations with postsynaptic specialization (GO:0099572), postsynaptic density (GO:0014069), asymmetric synapse (GO:0032279), synaptic vesicle (GO:0008021), neuron to neuron synapse (GO:0098984), exocytic vesicle (GO:0070382), cell cortex (GO:0005938), neuron projection cytoplasm (GO:0120111), synaptic vesicle membrane (GO:0030672), and exocytic vesicle membrane (GO:0099501) were observed (Figures 8g). Proteins relatively enriched in the F2^+^ were prominently associated with mitochondria i.e., inner mitochondrial membrane protein complex (GO:0098800), mitochondrial inner membrane (GO:0005743), mitochondrial protein-containing complex (GO:0098798), respiratory chain complex (GO:0098803), mitochondrial respirasome (GO:0005746), respirasome (GO:0070469), oxidoreductase complex (GO:1990204), mitochondrial respiratory chain complex I (GO:0005747), NADH dehydrogenase complex (GO:0030964), and respiratory chain complex I (GO:0045271) (Figures 8h).

To further validate our method and assess the concordance with known EV-associated proteins, we compared the proteins detected in the F1 and F2 BDEVs samples (both +/− and human/mouse-derived) with protein entries of the Vesiclepedia database (Kalra *et al*, 2012; Pathan *et al*, 2019). For mouse BDEVs, the protein make-up (both F1 and F2) showed around 90% overlap (i.e. F1^−^: 91.69%; F1^+^: 89.89%; F2^−^: 92.24%; and F2^+^: 90.50%), with the Vesiclepedia database protein entries (Supplementary Figure S8a). Similarly, for human BDEVs, we found that over 90% of the proteins detected in F1^−^ (91.83%) and F1^+^ (90.89%) as well as F2^−^ (91.82%) and F2^+^ (91.30%), have been previously reported to be associated with EVs (Supplementary Figure S8b).

### The mRNA analysis showed no differential expression comparing protocols, indicating a comparable mRNA BDEV content

With the previous experiments, we demonstrated that the isolated BDEV populations with and without collagenase were similar in terms of protein composition. To investigate similarities/differences among the BDEV preparations further, we then employed the Nanostring nCounter® Neurodegeneration panel to assess mRNA composition using the same protocol as previously described (Bub et al., 2022). For the analysis, we pooled F1 and F2 human BDEVs in both instances, isolated with (0.5 mg/mL) and without collagenase. Heatmaps and correlation plot analysis show no significant differences in mRNA expression profiles between collagenase-free and collagenase-based BDEVs since they did not cluster following the procedure used to isolate them (Figures 9a and 9b). The PCA plot indicates an overall similarity in the mRNA expression profiles of the data, although one of the collagenase-free samples was considered an outlier (Figure 9c). In essence, only two mRNA transcripts, *INA* and *SYT1*, were overrepresented, while three transcripts, *MOG, RING1,* and *PARP1*, were found to be deficient in the BDEV^−^, as depicted by the volcano plot (Figure 9d). In sum, these mRNA profiles further validate the similarities of the BDEV content isolated using both methods.

**Figure 9:**
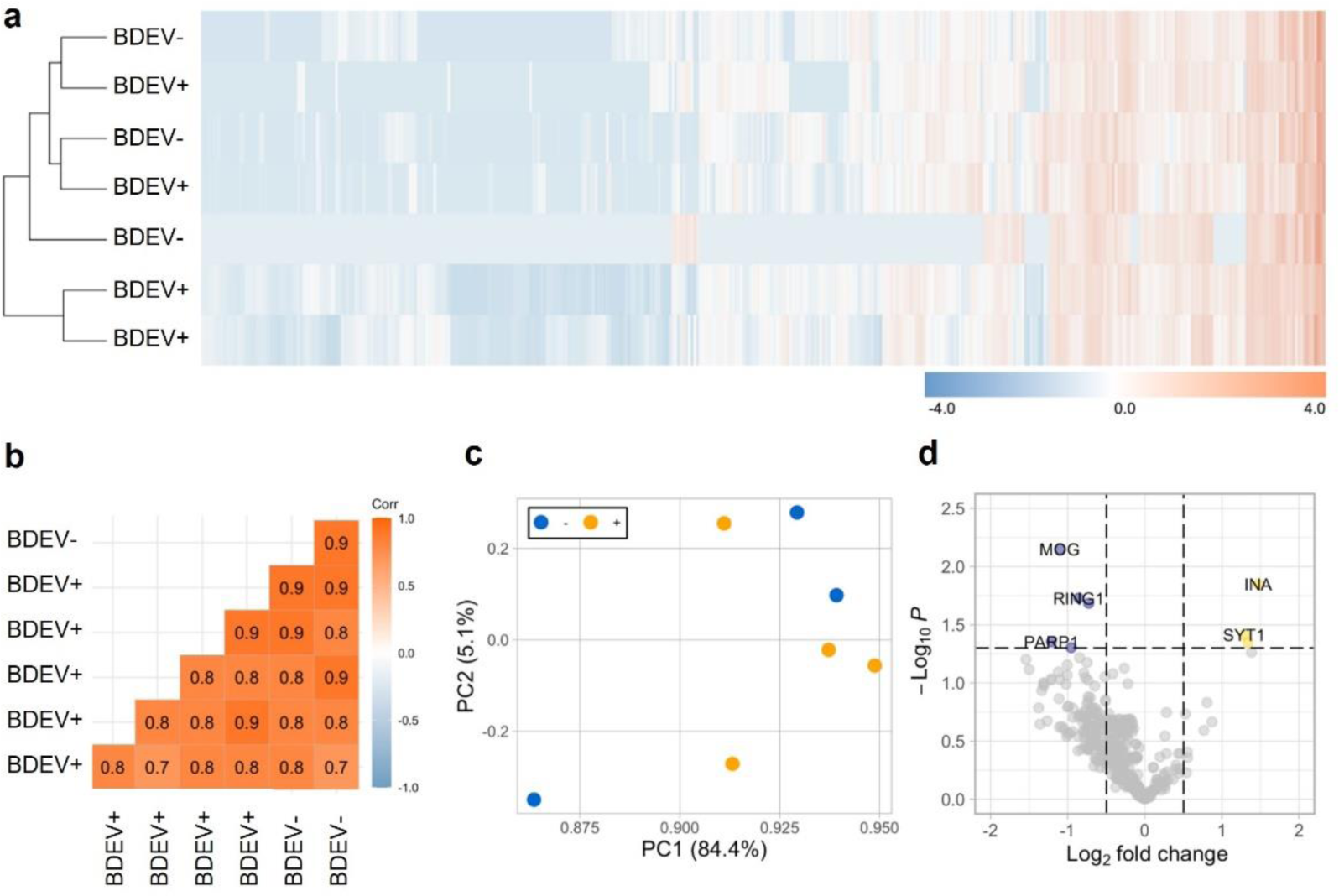
mRNA composition of the BDEVs isolated from human tissues displayed no differences between the collagenase-free and collagenase-based isolation protocols. (a) Heatmap displaying mRNA expressions (z-score) of BDEV^−^ (n = 3) and of BDEV^+^ (n = 4). (b) Correlation plot highlighting the similarities between collagenase-free and collagenase-based samples. The color key represents the variation in Pearson’s correlation coefficient values. (c) Scatter plots showing principal component analysis (PCA) of mRNA expression profiles of BDEV^−^ and BDEV^+^ EVs. (d) Volcano plot highlighting the downregulated (shown in blue) and upregulated (shown in yellow) mRNAs in the BDEV preparations. The x-axis intercepts denote the thresholds for expression changes, established at −1.5 fold for down-regulation and +1.5 fold for up-regulation. Meanwhile, the y-axis intercept signifies the threshold set for a *p*-value of 0.05.

## Discussion

In this study, we further investigated our observation that collagenase-based BDEV isolation leads to artificial cleavage of PrP^C^ (Brenna *et al*, 2020), to other EV markers such as CD81 and Flotillin-1. Our experiments involving collagenase treatment on mouse and human brain homogenates, on BDEVs, and on N2a-EVs confirm that the altered protein cleavage can be directly attributed to collagenase treatment. Our approach was to use a recently published protocol to isolate BDEVs from tissue with some modifications (Crescitelli *et al*, 2021) and compare samples isolated with a lower amount of enzymatic digestion with collagenase D (0.5 mg/mL) or no enzymatic digestion at all. We could demonstrate that without using collagenase D, we could still isolate BDEVs with high yield and purity, with the advantage of preserving protein integrity on the membrane. Our proteomic and mRNA expression analysis further validated that the new approach yields the same BDEV population as with the low collagenase-based isolation.

One of the initial interactions that EVs engage in after biogenesis is with the extracellular matrix (ECM), where EVs gain additional surface proteins (the so-called corona) through various chemical and biophysical mechanisms, including hydrogen and covalent bonding (Debnath *et al*, 2023). This corona bestows EVs with different biological properties (Tóth *et al*, 2021; Wolf *et al*, 2022). One of the key considerations in EV isolation from tissue is the trade-off between efficient releasing EVs from the ECM while avoiding intense cell and EV breakage, which can contaminate the EV preparations with cell debris and intracellular vesicles, and vesicle-like structures such as endosomes, lysosomes, Golgi-derived vesicles, and nuclear fractions. For this reason, it is crucial not to use a harsh tissue homogenization process and a specific separation procedure to separate the EVs from the cell debris (e.g., density-based gradients and differential centrifugation). The EV isolation from brain tissues includes the use of proteases of different types to help in this process, such as collagenases and papain (Vella *et al*, 2017; Cheng *et al*, 2018; Su *et al*, 2021; D’Acunzo *et al*, 2021), to break down the associations of EVs with the ECM and release them (Debnath *et al*, 2023). However, these proteases, due to their exo-proteolytic activity, could also induce unwanted protein cleavages (Duarte *et al*, 2016), leading to unwanted degradation or trimming of EVs-associated proteins possibly affecting both their experimental analysis and physiological behavior.

Enzymatic dissociation is the method of choice for preparing primary cultures from tissue and has been studied in detail to ensure the perfect balance between the optimal isolation of cells and minimal artificial modification. For instance, previous studies have explored the effects of such enzymatic-based isolation protocols in various cell types and tissues: satellite stem cells from muscle tissue (Miersch *et al*, 2018), mesenchymal stem/stromal cell isolation from umbilical cord tissue (Taghizadeh *et al*, 2018), adipose-derived stem cells (Koellensperger *et al*, 2022), chondrocytes from cartilage (Hidvegi *et al*, 2006), T cells from spleen and lung tissue (Liu *et al*, 2018) and neurons and glial cells from brain cortex (Panchision et al, 2007; Mattei et al, 2020; Hu et al, 2022). Similarly, O’Flanagan *et al*. reported that the collagenase digestion for dissociating solid tumor tissues triggers a stress response that alters the transcriptome of the isolated cancer cells (O’Flanagan *et al*, 2019). In line with this, Mattei *et al*. described that the enzymatic digestion of fresh brain tissue induces critical and consistent alterations in the proteome and transcriptome of neuronal and glial cells (Mattei *et al*, 2020). These studies highlight the impact of diverse enzyme-based cell isolation protocols on surface proteins that critically affect downstream flow cytometry analysis (Taghizadeh *et al*, 2018; Liu *et al*, 2018; O’Flanagan *et al*, 2019; Mattei *et al*, 2020). This relates to our findings concerning EV-membrane proteins such as CD81 and PrP^C^ and demonstrates the relevance of enzyme-free isolation.

Among the proteins that are altered with the collagenase D treatment we found CD81, which shows two bands after enzyme-based isolation, which disappear when using the enzyme-free protocol. The concrete role of CD81 on EVs is still not fully understood, and it seems to be dependent on the cell origin of the vesicles. However, CD81 has been related to crucial intracellular processes such as immune activation (e.g., CD81 forms a complex with the receptor CD9 that is critical for B cell development and activation) (Susa *et al*, 2021; Toribio & Yáñez-Mó, 2022; Fan *et al*, 2023), functions that could be affected by its pruning when isolating the EVs with enzymes. Similarly, the enzymatic digestion could also explain the double band observed for Flotillin-1, although we did not observe this effect in the digested brain homogenate. This may suggest a lower affinity of the collagenase D to digest this protein in a highly concentrated sample as the brain homogenates compared to the EV fraction, where Flotillin-1 is highly enriched.

Our focus of research is on the role of PrP^C^ on EVs in the context of neurodegenerative diseases (Falker *et al*, 2016; Heisler *et al*, 2018). Therefore, it was crucial for us to study how to preserve PrP^C^’s integrity in BDEVs. PrP^C^ is a GPI-anchored membrane protein, but it is also found in the extracellular space as soluble forms and on EVs (Matamoros-Angles *et al*, 2023). The first evidence of the PrP^C^’s presence on EVs was described almost two decades ago (Fevrier *et al*, 2004). After that, several publications have shown the important role of the EV-PrP^C^ in many functions related to brain pathophysiology, such as neuroprotection after ischemic conditions (Guitart *et al*, 2016), lysosomal-exosomal trafficking in neurons (Heisler *et al*, 2018), EV uptake (Brenna *et al*, 2020; D’Arrigo *et al*, 2021), and neurite outgrowth (Gonias *et al*, 2022). PrP^C^ experiences endogenous post-translational modifications and cleavages that generate several physiological PrP isoforms, both, membrane-anchored and released to the extracellular space that exert different functions. The two most frequent processes are the α-cleavage, produced by a still unknown protease, in the central part of the protein, generating two structurally different parts, and the shedding of PrP^C^, which is mediated by ADAM10 (see (Linsenmeier *et al*, 2017; Mohammadi *et al*, 2023). The enzymatic digestion induced by the collagenase in brain homogenates is similar but not identical to the naturally occurring alpha-cleavage since an additional larger fragment is formed in the case of collagenase treatment. However, we cannot exclude that this observed additional fragment is an intermediate form generated in a two-step process, and, therefore. The latter increases the α-cleavage-like processing of PrP^C^ on EVs (therefore increasing the amount of C1 fragment attached to the EVs, although they are already physiologically enriched in PrP-C1 as shown (Brenna *et al*, 2020). Moreover, this artificial modification observed here would induce the release of the N-terminal part of PrP^C^, which typically interacts with diverse ligands, potentially modulating their functions (see (Shafiq *et al*, 2022)), such as synaptic receptors in epilepsy (Carulla *et al*, 2011), the beta-amyloid in Alzheimer’s disease (Laurén *et al*, 2009; Resenberger *et al*, 2011; Falker *et al*, 2016), the α-synuclein in Parkinson models (Urrea *et al*, 2018) or the misfolded PrP in Prion diseases (Turnbaugh *et al*, 2011). Moreover, the lack of this N-terminal part could also affect the interaction of EVs with recipient cells and their uptake since it has been shown that PrP^C^ expression modulates the EV uptake (Brenna *et al*, 2020). Therefore, any functional analysis aiming to study PrP^C^ on EVs needs to ensure the physiological composition of the various PrP forms, avoiding any undesired proteolytic processes to preserve their function completely and prevent any bias in the downstream experiments.

Surprisingly, we observed a GM130 positive signal when BDEVs were isolated without enzymes. Our results with brain homogenates treated with collagenases (III and D) demonstrate that the GM130 “disappearance” is directly connected to collagenase digestion. Moreover, in our isolation controls, the 10K pellet (containing large EVs) and the pre-gradient BDEVs, already show a negative signal, highlighting that the digestion is previous to the small BDEV purification. However, GM130 was not detectable in the isolation of EVs from cell culture media of N2a cells. One possibility is that, on some occasions, GM130 is present in some BDEV subpopulations. However, our proteomic data contradicted this idea as it showed that the EV populations that we isolated with and without collagenase are quite similar. Another possibility is that GM130 is released during the tissue preparation from disrupted cells, sticking to the EV corona and that collagenase is degrading this extracellularly released GM130. The fact that we do not see it in the isolation of EVs from media (without significant cell disruption and not using collagenase) or in the isolation of EVs from blood (data not shown), would speak more in this direction. However, given the fact that the presence of GM130 has been described in at least 20 entries of mass spectrometry data in Vesiclepedia (Kalra *et al*, 2012; Pathan *et al*, 2019), we cannot completely rule out the possibility that, in some instances, GM130 can be present in EVs.

In conclusion, our results show that collagenase D treatment leads to the degradation of specific membrane proteins in BDEVs and suggest that caution should be taken into account when using enzymes for EV isolation from complex tissue such as the brain. Overall, BDEVs isolated without collagenase show a comparable purity compared to the ones isolated with a lower amount of collagenase. Moreover, isolation approaches obtained a similar EV population enrichment with just some minor proteomic and mRNA differences. The relevant advantage of this approach is that some key proteins at the EV membrane are not artificially cleaved, allowing more physiological functional EV studies. However, further studies are needed to fully characterize the isolated BDEVs to understand the biological implications of our findings and to explore their potential applications in various disease contexts.

## Acknowledgments

The authors would like to thank Prof. Dr. Lucie Carrier, Saskia Schlossarek, and Elisabeth Krämer from the Nanostring Core Facility of the University Medical Center Hamburg-Eppendorf (UKE) for their help and sample analysis with the nCounter^®^ panel. Figures 1 and 2 were created with BioRender.com.

## Authors’ contributions

AM-A, EK, and MS performed most of the experiments and data collection and wrote the original draft. MS did the bioinformatic analysis of all Proteomics and nCounter^®^ data. The BDEV isolation, protocol optimization, and BDEV characterization, N2a-derived experiments were done mainly by AM-A and EK, with the technical help of FS, BM, and MS. HCA, SB, and BP provided advice and expertise about the BDEV isolation protocol and derived techniques, being indispensable for starting the project. The TEM was performed by MSc and CS, together with the hands-on of EK, AM-A, and MS. IF provided the human samples included in this project. BS, HV, and HS performed the biochemical processing of the proteomic analysis and provided the proteomic raw data. AM-A, MS, and MG designed and supervised the project. BP, MK, SB, HCA, BM, and MG provided critical feedback during the entire project development and edited the manuscript. All authors reviewed the manuscript and read and approved the final version.

## Declaration of Interest Statement

The authors declare no conflict of interest.

## Funding

AM-A’s contributions are supported by the European Union’s Horizon 2020 research and innovation program through the Marie Skłodowska-Curie grant agreement N° 101030402. FS is supported by the China Scholarship Council (No. 202108080249). Additionally, MS and MG express their gratitude for financial support from various sources: the Joachim Herz Stiftung in Hamburg, Germany; the PIER Hamburg/Boston seed grant (PHM-2019-03) and PIER seed grant (PIF 2020-10) from the University of Hamburg, Germany, for MS; and the Forschungsförderungsfonds der Medizinischen Fakultät grant (NWF-20/10) from the University Medical Center Hamburg-Eppendorf, Germany, for MS. HCA was supported by the CJD Foundation (USA) and the Alzheimer Forschung Initiative e.V. (AFI, Germany).

## Notes

### Competing Interest Statement

The authors have declared no competing interest.

