## Supplementary Figures for "Efficient enzyme-free isolation of brain-derived extracellular vesicles"

**Supplementary Figures for Matamoros-Angles and Karadjuzovic *et al.***

**Supplementary Figure S1: Collagenase treatment effect on human brain tissue**

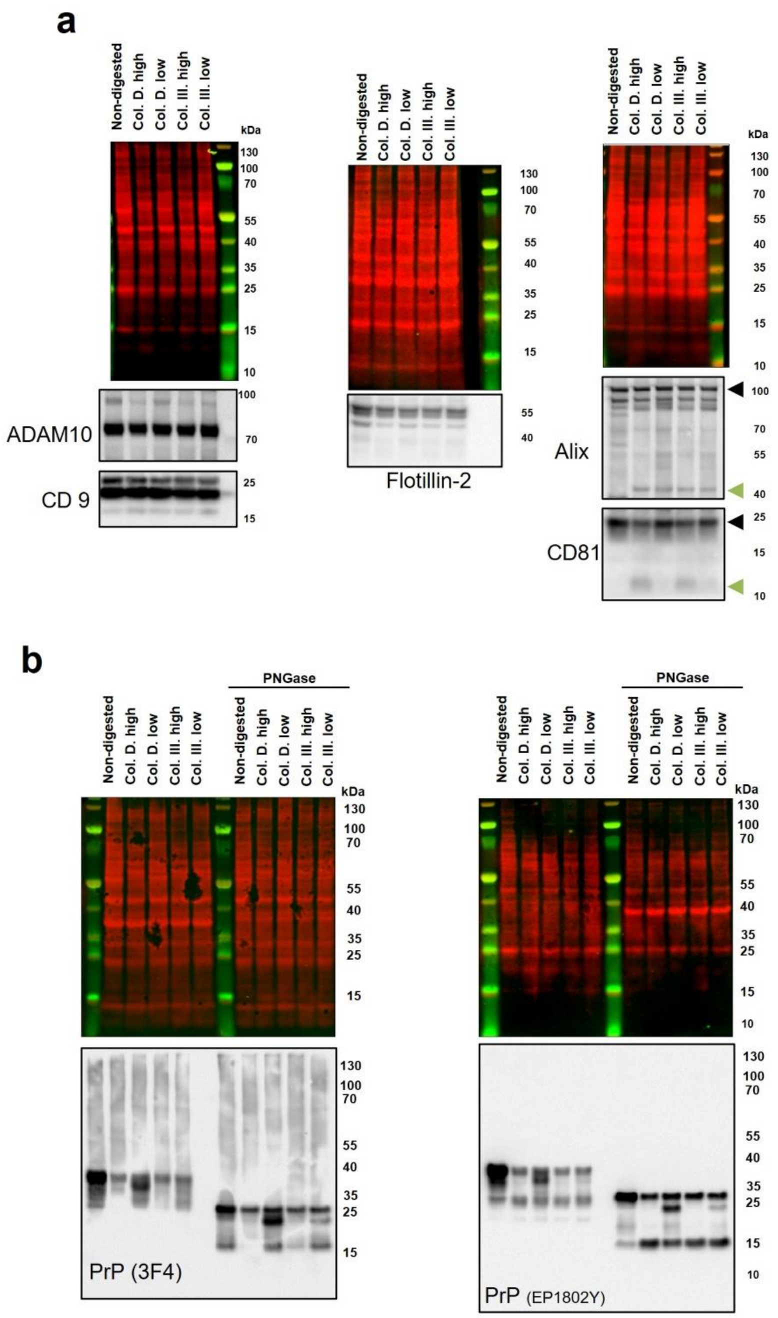


**Supplementary Figure S1: Collagenase treatment effect on human brain tissue.**

(a) Western blotting of human brain homogenates after the digestion with collagenase D and collagenase III. The same samples were loaded three times. CD81 and Alix presented an altered pattern but not CD9, ADAM10, or Flotillin-2. In (b), the same samples with PNGase treatment were loaded 2 times. $PrP^C$ was detected with two antibodies, one C-terminal (EP1802Y) and the other more N-terminal (3F4). The N-glycan removal showed exacerbated PrP-C1-like, PrP-N1-like forms, and an artificial shorter full-length $PrP^C$, as was previously observed with mouse brain after collagenase digestion.

Supplementary Figure S2: Images of the uncropped blots and total protein stainings from Figure 1.

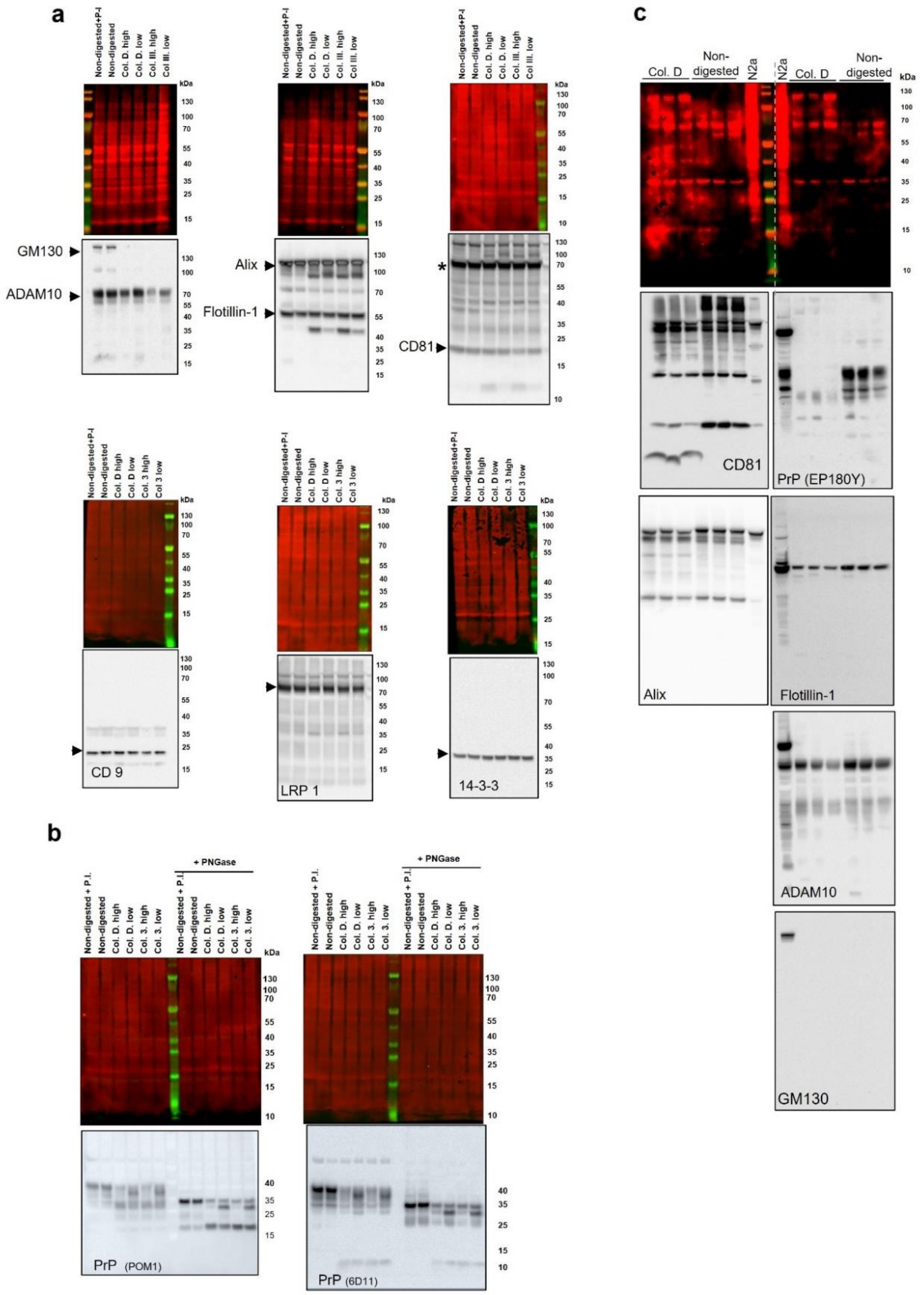


**Supplementary Figure S2: Images of the uncropped blots and total protein stainings from Figure 1.** (a) The same samples were loaded six times. Complete blots and total protein stainings of Figure 1b. GM130 was detected, and later, ADAM10 was reprobed. Alix was detected, and Flotillin-1 was reprobed. SDHA (marked as *) was detected (but not included in the final figure of the manuscript), and CD81 was reprobed. CD9, LRP1, and 14-3-3 were detected separately. (b) Complete blots and total protein staining of the western blotting of Figure 1c. Finally, in (c), the complete blots and total protein staining of Figure 1h are shown. The same samples were loaded two times. Alix was detected, and CD81 was later reprobed. On the other membrane, $PrP^C$, Flotillin-1, ADAM10 and GM130 were detected sequentially with a protein stripping in between.

**Supplementary Figure S3: EV expression markers in mouse tissue BDEV isolation controls and fractions: 10,000*xg* pellet, pregradient BDEVs, F3 and F4.**

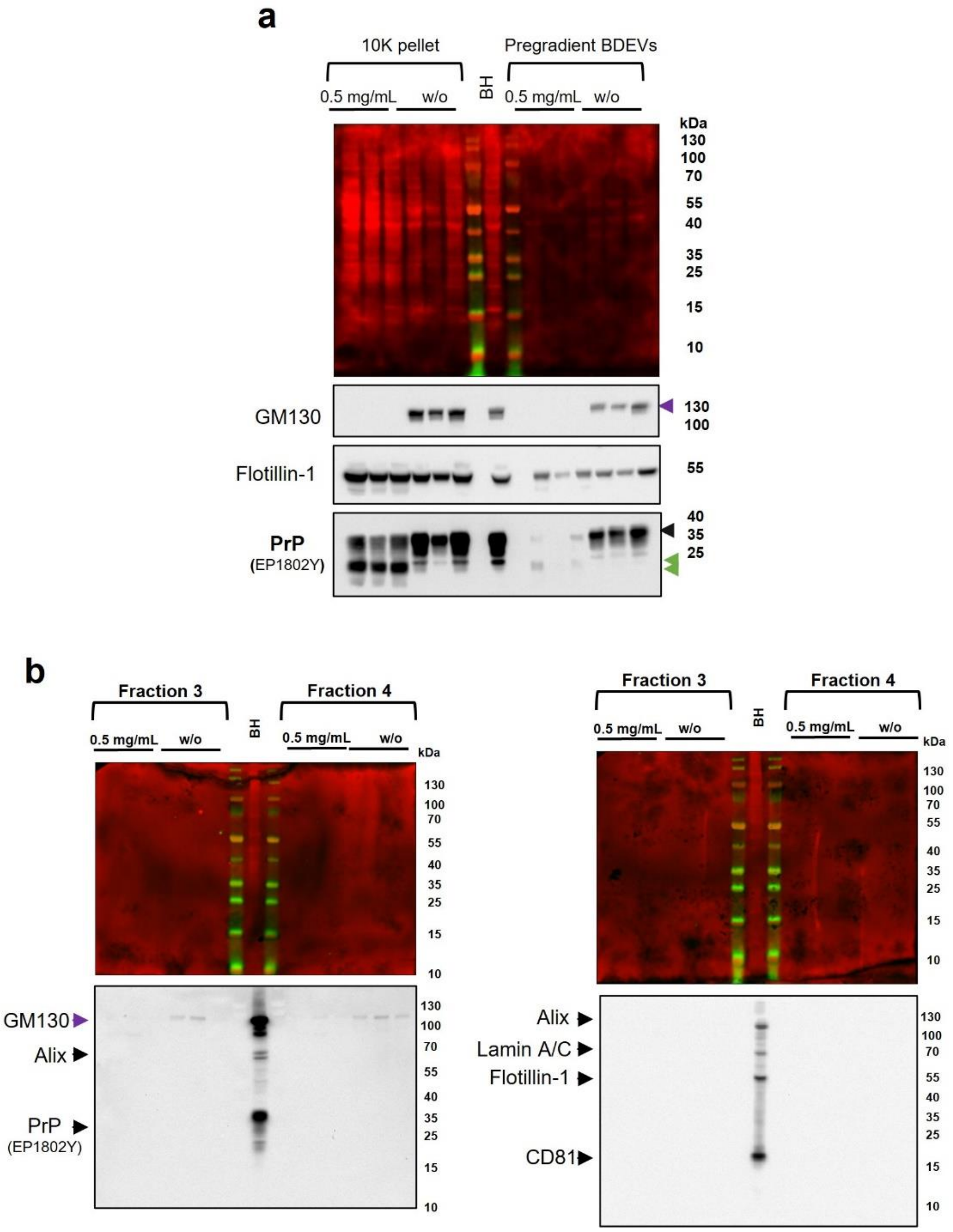


**Supplementary Figure S3: EV expression markers in mouse tissue BDEV isolation controls and fractions: 10,000*xg* pellet, pre-gradient BDEVs, F3 and F4.**

Expression of EV positive markers (Alix, CD81, and Flotillin-1), EV negative markers (Lamin A/C and GM130), and $PrP^C$ (EP1802Y) in the BDEV isolation control samples obtained during the isolation of the BDEVs shown in Figure 3. The same samples were loaded two times. In (a), the 10,000*xg* pellet (10K) and the pre-gradient BDEVs showed a GM130 total disappearance when collagenase was used, and $PrP^C$ displayed the cleavage pattern observed in the BDEVs. In (b), the GM130 signal was detectable in fraction 3 (F3) and fraction 4 (F4) in the samples obtained with 0 mg/mL of collagenase, but there is no expression of any other marker. The total protein staining also shows no protein expression in the samples, but it does in the total mouse brain homogenate (BH) used as a loading control.

**Supplementary Figure S4: EV expression markers in human tissue BDEV isolation controls and fractions: 10,000*xg* pellet, pregradient BDEVs, F3 and F4.**

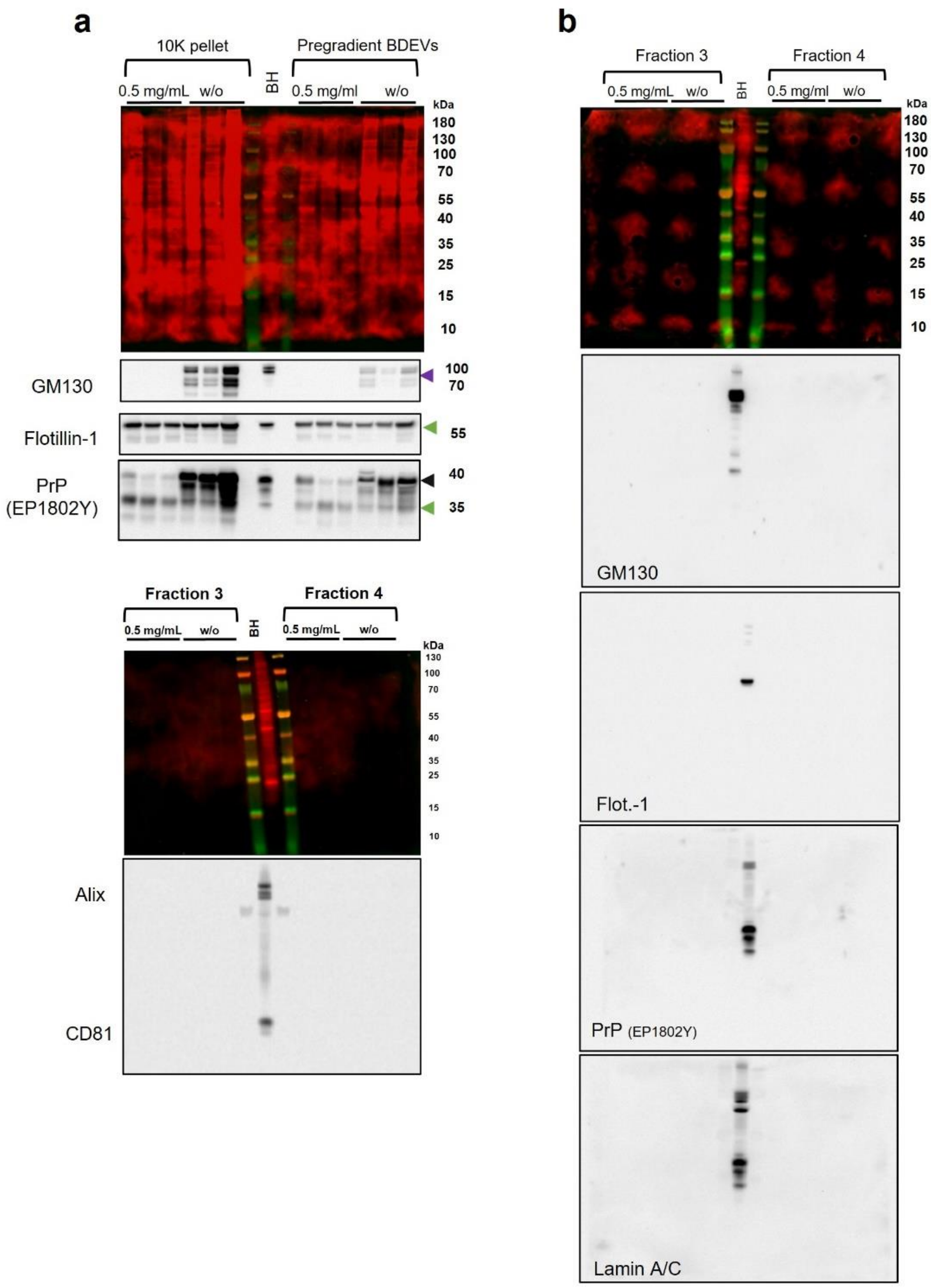


**Supplementary Figure S4: EV expression markers in human-BDEV isolation controls and fractions: 10,000*xg* pellet, pre-gradient BDEVs, F3 and F4.**

Expression of EV positive markers (Alix, CD81, and Flotillin-1), EV negative markers (Lamin A/C and GM130), and $PrP^C$ (EP1802Y) in the BDEV isolation control samples obtained during the isolation of the BDEVs shown in Figure 4. The same samples were loaded two times. In (a), the 10,000*xg* pellet (10K) and the pre-gradient BDEVs showed a GM130 total disappearance when collagenase was used, and $PrP^C$ displays the cleavage pattern observed in the BDEVs. In (b), fraction 3 (F3) and fraction 4 (F4) showed no expression of any marker. The total protein staining also shows no protein expression in the samples, but it does in the total human brain homogenate (BH) used as a loading control.

Supplementary Figure S5: Images of the uncropped blots and total protein stainings from Figure 3 and Figure 4.

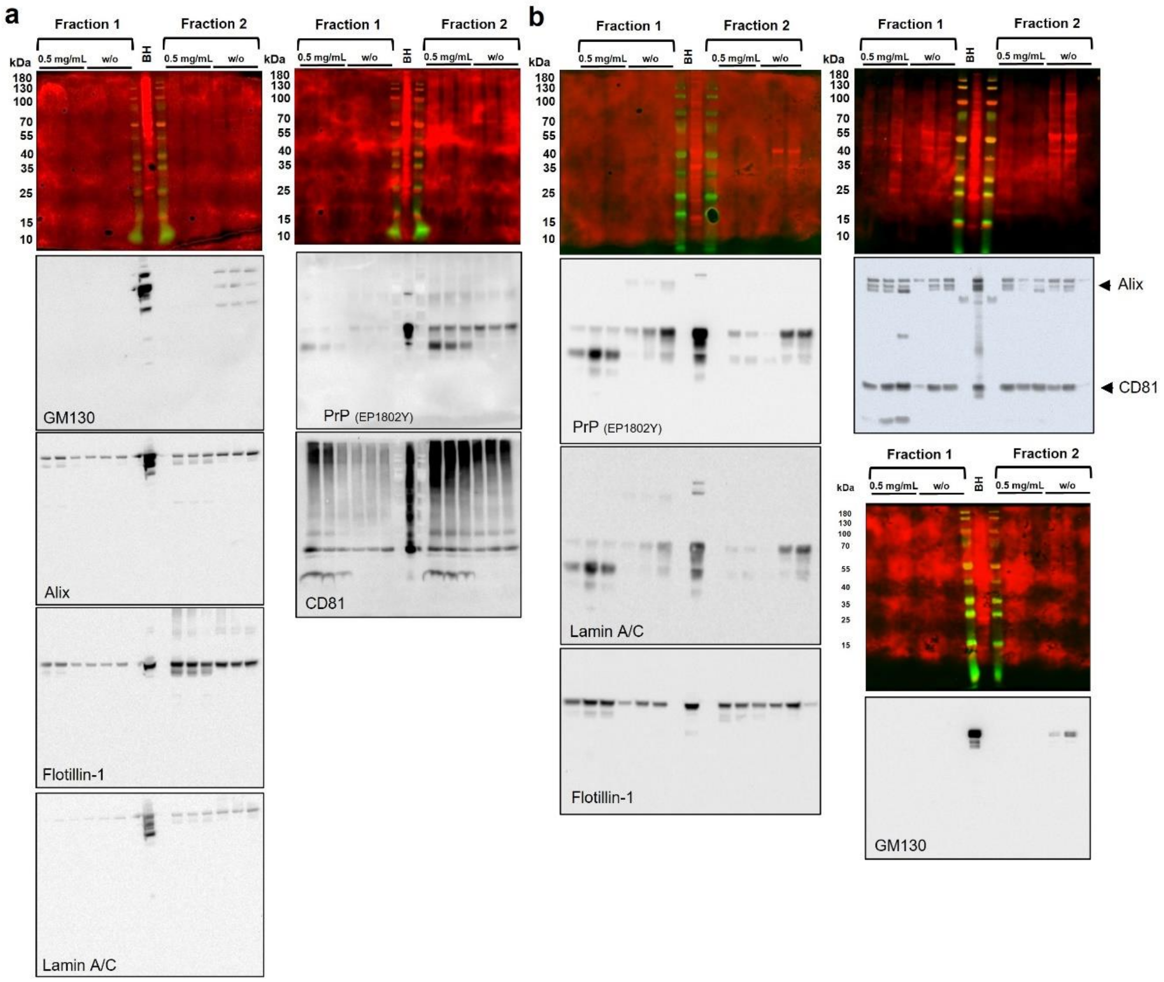


**Supplementary Figure S5: Images of the uncropped blots and total protein stainings from Figure 3 and Figure 4.**

(a) The same samples were loaded twice with their corresponding uncropped films of Figure 3a. GM130 was initially detected, followed by an intercalated stripping, Alix, Flotillin-1, and Lamin A/C were detected. On the other membrane, $PrP^C$ (using EP1802Y antibody) was detected, and then CD81 was reprobed. (b) The same samples were loaded three times with their corresponding uncropped films in Figure 4a. $PrP^C$ was initially detected in the first membrane, and later, after an intercalated stripping, Flotillin-1 and Lamin A/C were detected. On the other membrane, first, Alix was detected, and then CD81 was reprobed. Finally, GM130 was detected alone on the last membrane.

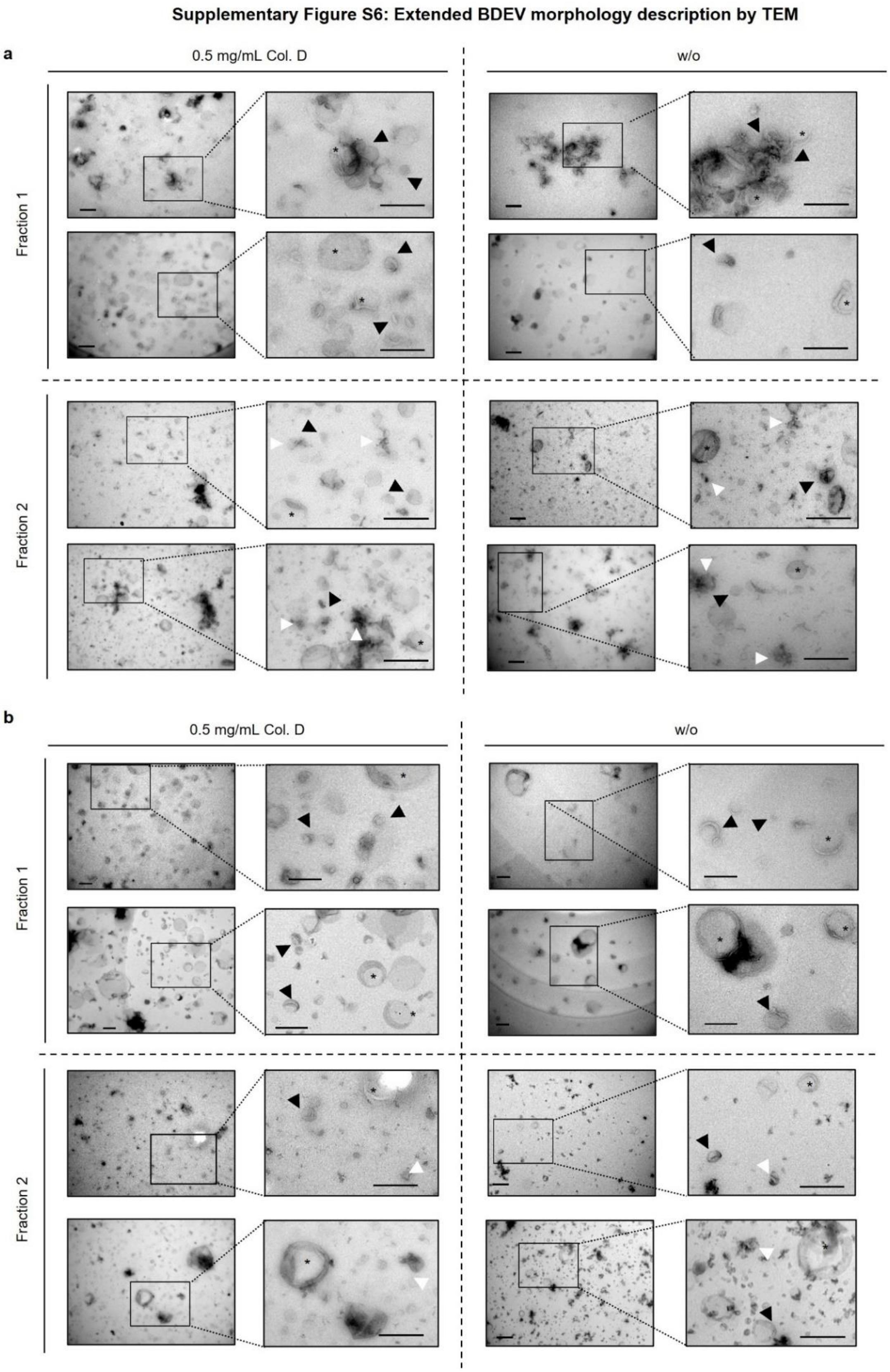


**Supplementary Figure S6: Extended BDEVs morphology description by TEM.**

TEM images of mouse (a) and human (b) BDEVs obtained with (0.5 mg/mL) or without (w/o) collagenase. Detailed membranous BDEV structures are observed in all the samples (arrowheads), and also large vesicles folded (asterisk). In both cases, as expected, the F2 shows more debris and aggregated structures. Scale bars = 500 nm.

**Supplementary Figure S7: Collagenase-based (2mg/ml) BDEV isolation produces aberrant cleavage of EV proteins but not significantly higher yield. Uncropped blot images and total protein staining along with BDEVs F3-F4 EVs marker expression.**

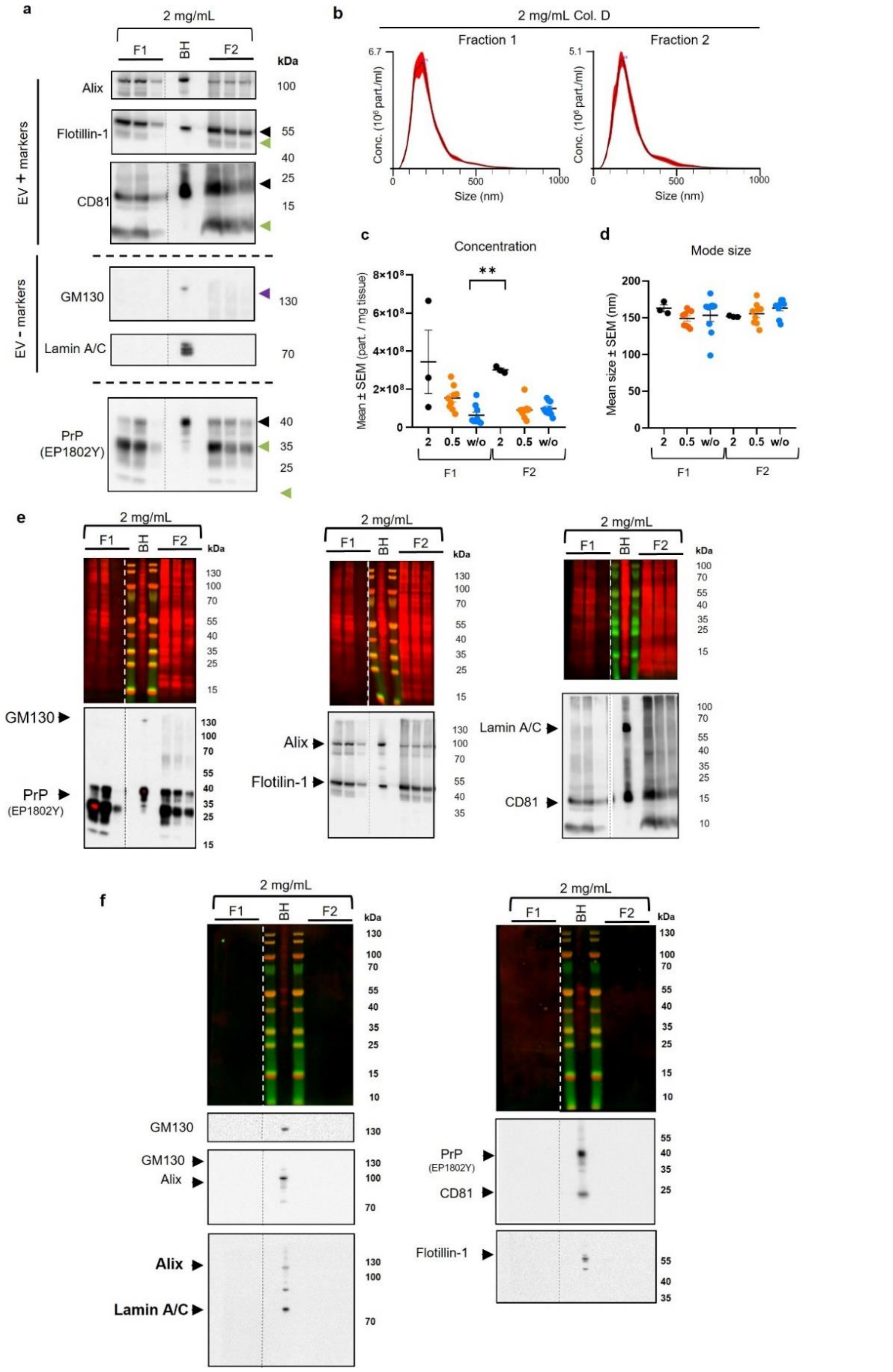


**Supplementary Figure S7: Collagenase-based (2 mg/mL) BDEV isolation produces aberrant cleavage of EV proteins but not significantly higher yield. Uncropped blot images and total protein staining along with BDEVs F3-F4 EVs marker expression.**

(a) Representative western blots of F1 and F2 BDEVs samples isolated from mouse brain tissue, with 2 mg/mL of collagenase D (*n* = 3 preparations/condition), for PrP$^{C}$, the EV positive markers Alix, Flotillin-1, and CD81 and the Golgi and nucleus markers, GM130 and Lamin A/C, as EV negative markers. The green arrowheads indicate the artificial cleavage observed in the proteins $PrP^C$, Flotillin-1 and CD81, and the purple one indicates the unexpected presence of GM130. A total mouse brain homogenate (BH) was used as a loading control. (b) Representative size distribution graphs from NTA analysis of F1 and F2 BDEVs. The concentration of particles per mg of initial tissue (c) and the size (d) of the BDEVs obtained with this protocol is not different in any fraction compared with the ones isolated with 0.5 mg/mL collagenase D or without (w/o) any collagenase (data from Figure 3). Just the $F1^-$ has significantly less concentration than the $F2^{2+}$ (**$p$ = 0.005, Kruskal-Wallis test). Data are presented as mean ± S.E.M. in (c) and (d). (e) Uncropped images of the blots shown in panel (a) with their respective total protein stainings. To be able to detect a greater number of markers and proteins by western blot, the same set of samples were loaded three times. The detection sequence involved GM130 and subsequent reprobing for $PrP^C$ on the first membrane. On the second membrane, Flotillin-1 was detected, and later, it was reprobed for Alix. Lastly, on the third membrane, Lamin A/C, was detected, and it was later reprobed with CD81. (f) Expression of EV positive markers (Alix, CD81, and Flotillin-1), EV negative markers (Lamin A/C and GM130), and PrP (EP1802Y) in the fraction 3 (F3) and fraction 4 (F4) of the BDEVs presented in panel (a). The total protein staining shows no protein content in the F3 and F4 samples; likewise, no EV marker was detectable in these samples, in contrast to the total mouse brain homogenate (BH), which was used as a loading control.

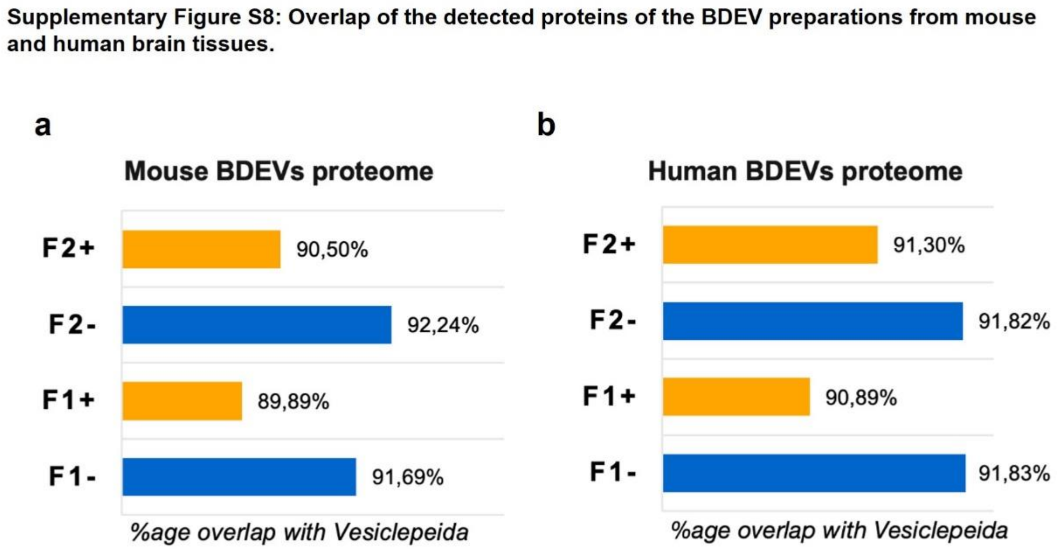


**Supplementary Figure S8: Overlap of the detected proteins of the BDEV preparations from mouse and human brain tissues.**

Proteins constituting the EV preparations were checked for their presence in the Vesiclepedia database using the FunRich enrichment software. (a) Mouse BDEV samples obtained with collagenase-free and collagenase-assisted methods from both F1 and F2 showed higher overlaps for their constituting proteins, with the Vesiclepedia database around 90%. (b) Likewise, over 90% of proteins from F1 and F2 EVs from both preparations of human BDEV were found in the Vesiclepedia database.
